# Targeting cancer glycosylation repolarizes tumor-associated macrophages allowing effective immune checkpoint blockade

**DOI:** 10.1101/2021.04.11.439323

**Authors:** Michal A. Stanczak, Natalia Rodrigues Mantuano, Nicole Kirchhammer, David E. Sanin, Jinyu Wang, Marcel P. Trefny, Gianni Monaco, Melissa A. Gray, Adam Petrone, Abhishek S. Kashyap, Katharina Glatz, Benjamin Kasenda, Karl Normington, James Broderick, Li Peng, Oliver M.T. Pearce, Erika L. Pearce, Carolyn R. Bertozzi, Alfred Zippelius, Heinz Läubli

**Affiliations:** Department of Biomedicine, University Hospital and University of Basel, Switzerland; Max Planck Institute of Immunobiology and Epigenetics, Freiburg, Germany; Department of Chemistry, Stanford ChEM-H, and Howard Hughes Medical Institute, Stanford University, Stanford, CA, USA; Palleon Pharmaceuticals, Waltham, MA, USA; Institute of Pathology, University Hospital Basel, Switzerland; Division of Oncology, Department of Internal Medicine, University Hospital Basel, Switzerland; Centre for Tumour Microenvironment, Barts Cancer Institute, Queen Mary University, London, United Kingdom; The Bloomberg-Kimmel Institute for Cancer Immunotherapy at Johns Hopkins, Baltimore, MD 21287, USA

**Author notes:** Correspondence should be addressed to M.A.S. and H.L. These authors contributed equally to this work (Natalia Rodrigues Mantuano, Nicole Kirchhammer).

**Keywords:** sialic acid, Siglec, tumor immunology, tumor-associated macrophages, immune checkpoint, immunotherapy

## Abstract

Immune checkpoint blockade (ICB) has significantly improved the prognosis of cancer patients, but the majority experience limited benefit, evidencing the need for new therapeutic approaches. Upregulation of sialic acid-containing glycans, termed hypersialylation, is a common feature of cancer-associated glycosylation, driving disease progression and immune escape via the engagement of Siglec-receptors on tumor-infiltrating immune cells. Here, we show that tumor sialylation correlates with distinct immune states and reduced survival in human cancers. The targeted removal of Siglec-ligands in the tumor microenvironment, using an antibody-sialidase conjugate, enhances anti-tumor immunity and halts tumor progression in several mouse tumor models. Using single-cell RNA sequencing, we reveal desialylation mechanistically to repolarize tumor-associated macrophages (TAMs) and identify Siglec-E on TAMs as the main receptor for hypersialylation. Finally, we show genetic and therapeutic desialylation, as well as loss of Siglec-E, to synergize with ICB. Thus, therapeutic desialylation represents a novel immunotherapeutic approach, shaping macrophage phenotypes and augmenting the adaptive anti-tumor immune response.

## Introduction

Cancer immunotherapy using immune checkpoint blockade (ICB), including antibodies blocking CTLA-4 and PD-(L)1, has improved the outcomes of cancer patients, although overall only a minority benefit from the currently available ICB^1,2^. New target pathways are under investigation and combination approaches with CTLA-4- and PD-(L)1-blocking agents show promising preclinical and early clinical activity^3^.

The upregulation of sialic acid-containing glycans in the tumor microenvironment, termed tumor hypersialylation, contributes to the establishment of an immunosuppressive milieu and dampens anti-tumor immune responses via the engagement of immunomodulatory Siglecs expressed on tumor-infiltrating immune cells^4,5^. Recent work has suggested the sialoglycan–Siglec axis as a new immune checkpoint that can be targeted to drive innate and adaptive anti-tumor immunity^4,6-12^. However, given the existence of multiple Siglecs and their broad pattern of expression in the immune system, the exact mechanism remains unclear.

The expression of inhibitory CD33-related Siglecs, including human Siglec-7/-9 and murine Siglec-E, on tumor-associated macrophages (TAMs) was shown to support cancer progression by driving macrophage polarization towards the tumor-promoting M2 phenotype^4,9,13^. Similarly, NK cell-mediated killing of tumor cells can be blocked in a dose-dependent manner by the interactions between tumor sialoglycans and Siglec-7/-9 on human NK cells^7,11^. Recent work by us and others identified Siglec-9 as an inhibitory receptor expressed on tumor-infiltrating T cells in different cancers, including non-small cell lung cancer, epithelial ovarian cancer, colorectal cancer and melanoma^6,10^. We previously showed that genetic desialylation of tumor cells halts tumor growth and leads to improved anti-tumor immunity in mouse models^6^ and demonstrated that treatment with sialidase can achieve similar results^14^. While a growing body of evidence supports the potential of targeting tumor sialylation, the feasibility of therapeutic interventions and synergy with classical ICB remained to be demonstrated.

Here, we investigate the cellular and molecular mechanisms by which therapeutic desialylation augments anti-tumor immunity and halts tumor growth. We identify Siglec-E expression on TAMs to mediate the effects of desialylation and demonstrate synergism in combination with ICB.

## Results

### Tumor sialylation is associated with immune suppression and reduced survival in cancer patients

As increased tumor sialylation has previously been linked with immune suppression^15,16^, we wanted to test if the expression of sialic acid-modifying enzymes is associated with distinct immune states in human cancer. To that end, we assembled a set of genes encoding for proteins involved in sialoglycan biosynthesis, each of which was then correlated and clustered with a previously published set of 3,021 immune genes using the gene expression data of all solid cancers from the Cancer Genome Atlas (TCGA) database. Based on the clustering, five sialylation gene sets were defined and tested for their association with patient survival (Fig. 1a, Extended Data Fig. 1a). This led to the identification of gene set 1, consisting of α-2,3- and α-2,6-sialyltransferases, which was significantly associated with regulation of immune function and reduced cytokine activity. Expression of gene set 1 strongly correlated with reduced survival of patients with several cancer types (Extended Data Fig. 1b), in particular patients with clear cell kidney cancer (KIRC, Fig. 1b) and squamous cell lung cancer (LUSC, Fig. 1c, Extended Data Fig. 1c). To corroborate this finding, we stained a tissue microarray of 75 primary human KIRC samples and corresponding healthy control tissues with a hexameric human Siglec-9 Fc protein, to quantify the expression of sialic acid-containing Siglec-ligands. Analysis of the linked survival data revealed a worse overall survival in KIRC patients with increasing Siglec-9 Fc-binding (Extended Data Fig. 1d, e), validating the use of gene set 1 expression as a proxy for hypersialylation and supporting the association with reduced survival.

**Figure 1.**
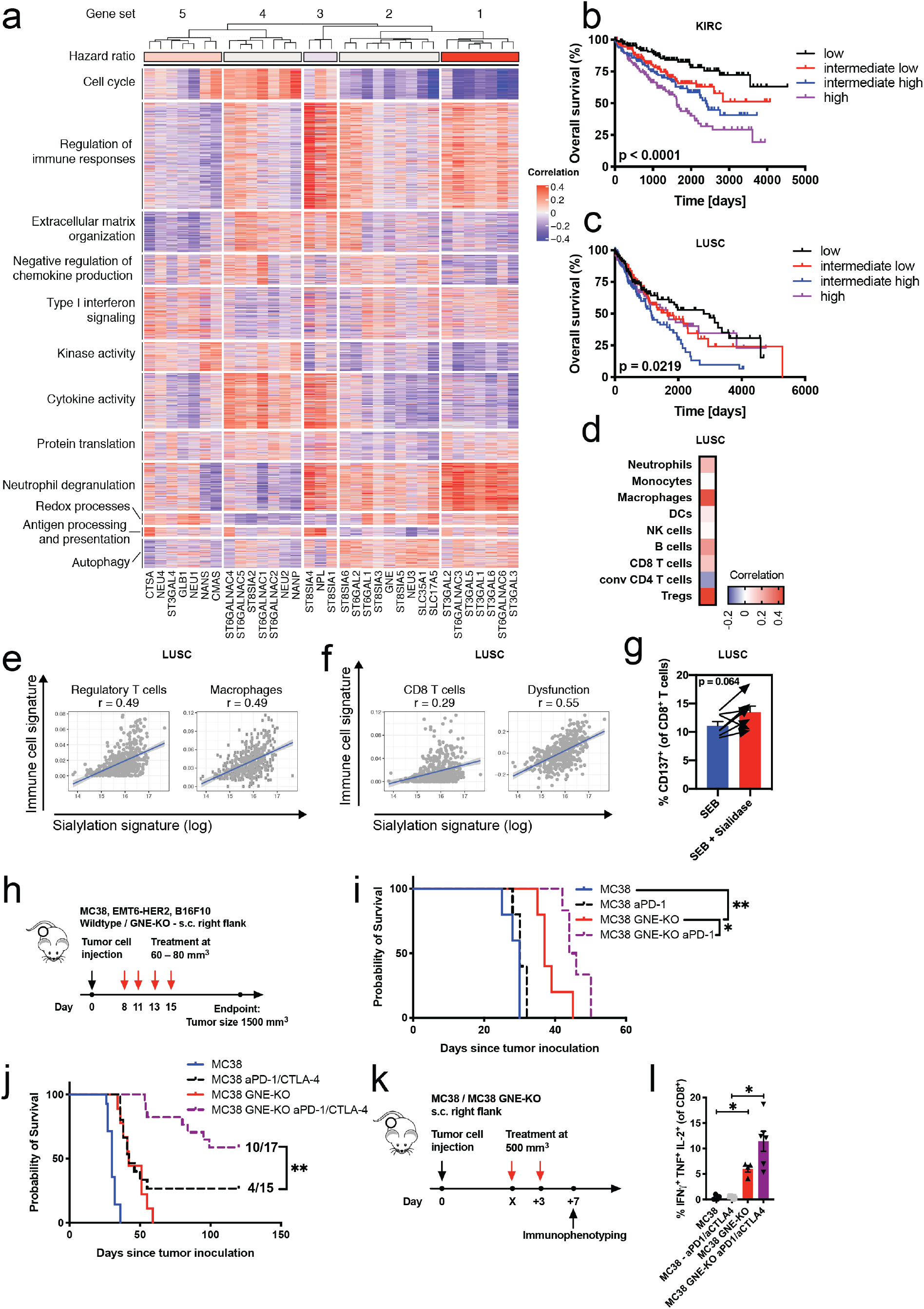
Tumor sialylation is associated with immune suppression and reduced survival in cancer patients. **a**, Clustering of correlations between the expression of sialic acid-modifying enzymes with the expression of immune genes in all solid cancers from the Cancer Genome Atlas (TCGA) database. **b**, Kaplan-Meier survival curve of patients with clear cell renal carcinoma (KIRC) or in **c**, squamous cell carcinoma of the lung (LUSC), divided into quartiles based on the low, intermediate–low, intermediate–high and high expression of gene set 1, from TCGA data. **d**, Correlations between gene set 1 expression and gene signatures of the listed tumor-infiltrating immune cell types in all LUSC patients, from TCGA data. r values are shown on a color scale, from blue to red. **e**, Dot plots displaying correlations between gene set 1 expression and signatures of regulatory T cells and tumor-associated macrophages. **f**, Dot plots displaying correlations between gene set 1 expression and signatures of CD8^+^ T cells and T cell dysfunction in cancer. **g**, Expression of CD137 (4-1BB) on CD8^+^ T cells in primary LUSC samples from individual patients after 48 h incubation with the superantigen SEB alone or in combination with *Vibrio cholerae* sialidase (n=7). **h**, Experimental design: Mice carrying subcutaneous or intramammary wildtype and GNE-KO tumors were treated i.p. with four doses of 10 mg/kg anti-PD-1 and/or anti-CTLA-4 antibodies, beginning at a tumor size of approximately 80 mm^3^. **i**, Effect of PD-1 blockade on the survival of mice bearing subcutaneous wildtype or GNE-KO MC38 tumors (n=5–6 mice per group). **j**, Effect of combined PD-1 and CTLA-4 blockade on the survival of mice bearing subcutaneous wildtype or GNE-KO MC38 tumors (n=14–17 mice per group). **k**, Experimental design: Mice carrying established (approx. 500 mm^3^) subcutaneous wildtype and GNE-KO MC38 tumors were treated i.p. with two doses of 10 mg/kg anti-PD-1 and anti-CTLA-4 antibodies. 7 days after the first treatment, tumors were resected and immunophenotyped. **l**, Frequency of IFNγ^+^TNF^+^IL2^+^ CD8^+^ T cells after restimulation in single-cell suspensions of MC38 wildtype or GNE-KO tumors treated with PD-1 and CTLA-4 blockade (n=4–6 mice per group). n indicates the number of biological replicates. Error bars represent the mean ± standard error of the mean (s.e.m.). Statistical analyses were performed using the log-rank (Mantel–Cox) test for the TCGA survival data or the Gehan-Wilcoxon test for the mouse survival data, followed by Bonferroni’s correction for multiple comparisons. A paired two-tailed Student’s *t*-test was used in Fig. 1g and a one-way ANOVA with post hoc Sidak’s test in Fig. 1l. * *P* ≤ 0.05, ** *P* ≤ 0.01, *** *P* ≤ 0.001, **** *P* ≤ 0.0001.

Then, we correlated the expression of gene set 1 with gene expression signatures of different tumor-infiltrating immune cell types in all LUSC patients and found the strongest positive correlations with immunosuppressive and tumor-promoting cell types (Fig. 1d), such as regulatory T cells (Tregs) and TAMs (Fig. 1e). In contrast, the correlation with conventional CD4^+^ T cells was significant but negative (Extended Data Fig. 1f) and no correlations were found with other myeloid cell types such as dendritic cells (DCs) or monocytes (Extended Data Fig. 1g). While a small positive correlation could be observed with CD8^+^ T cells, it was notably weaker than the correlation with a published signature of T cell dysfunction in cancer (Fig. 1f)^17^. To test this correlation between tumor sialylation and T cell dysfunction, we polyclonally stimulated T cells in primary tumor suspensions from LUSC patients using the superantigen Staphylococcal enterotoxin B (SEB), alone or in combination with sialidase. In line with our findings, sialidase treatment lead to an increase in T cell activation, measured as the expression of the activation marker CD137 (Fig. 1g). These data link tumor sialylation to specific changes in immune infiltration, predominantly by immunosuppressive and tumor-promoting cell types and reduced survival of patients.

To corroborate the association between tumor sialylation and T cell dysfunction, we used a mouse tumor model of genetic desialylation in combination with anti-PD-1 and/or anti-CTLA-4 ICB. We previously showed that growth of tumors lacking the rate-limiting enzyme for sialic acid biosynthesis, UDP-GlcNAc 2-epimerase (GNE, GNE-KO), was delayed and survival of mice prolonged^6^. We confirmed the delayed tumor growth of MC38 GNE-KO tumors and were able to show the delay to be dependent on CD8^+^ T cells, evidenced by its abrogation when CD8^+^-depleting antibodies were applied (Extended Data Fig. 1h, i). In order to test the effect of tumor sialylation on the ability of ICB to reinvigorate tumor-infiltrating CD8 T cells, we subcutaneously injected mice with wildtype or GNE-KO MC38 tumor cells and treated mice with palpable tumors with four doses of PD-1 blocking antibodies alone, or in combination with CTLA-4 blocking antibodies (Fig. 1h). Treatment of GNE-KO tumors with anti-PD-1 resulted in a significantly stronger reduction in tumor growth and prolonged survival of mice compared to treatment of wildtype tumors (Fig. 1i, Extended Data Fig. 1j). Application of both PD-1 and CTLA-4 blockade led to the rejection of wildtype tumors in 4 of 15 mice (27%), whereas the rejection rate was increased to 10 of 17 (59%) in mice bearing GNE-KO tumors (Fig.1 j, Extended Data Fig. 1k).

In order to exclude a cell line or mouse strain specific effect, we used the EMT6-HER2 mammary carcinoma cell line injected orthotopically into the mammary fat pads of female BALB/c mice. Again, we observed a delayed tumor growth of GNE-KO tumors compared to wildtype tumors and an increased rejection rate of GNE-KO tumors treated with anti-PD-1 antibodies compared to wildtype tumors (50% vs. 33%, Extended Data Fig. 1l, m). Additionally, we applied the poorly immunogenic and highly aggressive melanoma B16F10 model, again demonstrating that GNE-KO delayed tumor growth, prolonged survival and increased sensitivity to PD-1/CTLA-4 blockade, resulting in a more pronounced delay in tumor growth and a longer survival of mice bearing GNE-KO tumors (Extended Data Fig.1n, o). To functionally test the reinvigoration of CD8 T cell function by ICB, we treated mice carrying established (approx. 500 mm^3^) wildtype or GNE-KO MC38 tumors with two doses of anti-PD-1 and -CTLA-4 blocking antibodies and isolated tumor-infiltrating immune cells 7 days after the first treatment (Fig. 1k). T cells from the single-cell suspensions of those tumors were restimulated *in vitro* and intracellularly stained for the expression of CD8^+^ effector T cell cytokines. While ICB alone increased the frequencies of both IFNγ^+^ and IFNγ^+^TNF^+^ CD8^+^ T cells (Extended Data Fig. 1r), desialylation augmented T cell activation, in particular when assessing the frequencies of multifunctional IFNγ^+^TNF^+^IL-2^+^ CD8^+^ T cells, which were not induced by ICB alone (Fig. 1l). These findings support the association between tumor sialylation, immunosuppression and dysfunction of T cells and demonstrate increased reinvigoration of tumor-infiltrating T cells by ICB in desialylated tumors.

### Tumor-targeted sialidase effectively desialylates the tumor microenvironment

Next, we aimed to study the cellular and molecular mechanisms underlying sialic acid-mediated immune suppression in greater detail using therapeutic desialylation in different mouse tumor models. In order to achieve tumor-specific desialylation *in vivo*, we utilized an antibody-sialidase fusion protein to target the enzyme to the tumor microenvironment. Specifically, we used the FDA approved antibody trastuzumab, which recognizes the HER2 receptor. The trastuzumab-sialidase construct, termed E-301, was bioengineered on the Enzyme-Antibody-Glycan-Ligand-Editing platform (*EAGLE*, Palleon Pharmaceuticals, WO2019136167) by fusing two sialidase domains to the C-terminal Fc region of trastuzumab. As controls, we used unmodified trastuzumab, as well as an enzymatically inactive E-301 variant carrying two loss-of-function (LOF) mutations in the catalytic sites of its sialidases (E-301 LOF, Fig. 2a). We confirmed dose-dependent desialylation of HER2 expressing EMT6 mammary carcinoma cells (EMT6-HER2) by E-301 but not by trastuzumab or E-301 LOF (Fig. 2b, Extended Data Fig. 2a). To evaluate tumor cell desialylation *in vivo*, we established orthotopic EMT6-HER2 tumors in BALB/c mice until they reached a size of 400–500 mm^3^ and treated the mice systemically with a single intraperitoneally (i.p.) administered dose of 10 mg/kg E-301 or trastuzumab (Fig. 2c). We observed increased staining with peanut agglutinin (PNA), detecting galactosyl residues uncovered by desialylation, and decreased staining with *Maackia amurensis* lectin II (MAL II), binding to α-2,3-sialic acids, in the E-301-treated tumors compared with the trastuzumab-treated and untreated tumors. Desialylation was confirmed both by immunofluorescence (Fig. 2d, e) and by flow cytometry (Fig. 2f) and was most pronounced at 24 h, but still detectable at 72 h post injection (Extended Data Fig. 2b, c). Staining with a Siglec-E Fc fusion protein at 72 h confirmed loss of Siglec-ligands after E-301 treatment (Extended Data Fig. 2d). Flow cytometric staining for human Fc further confirmed successful penetration and targeting of E-301 into the tumor but also suggested an accelerated clearance of E-301 over time compared to trastuzumab (Fig. 2f, Extended Data Fig. 2e). Our results confirm that targeting of sialidase to tumor cells results in a strong but transient reduction of sialoglycan levels *in vivo*.

**Figure 2.**
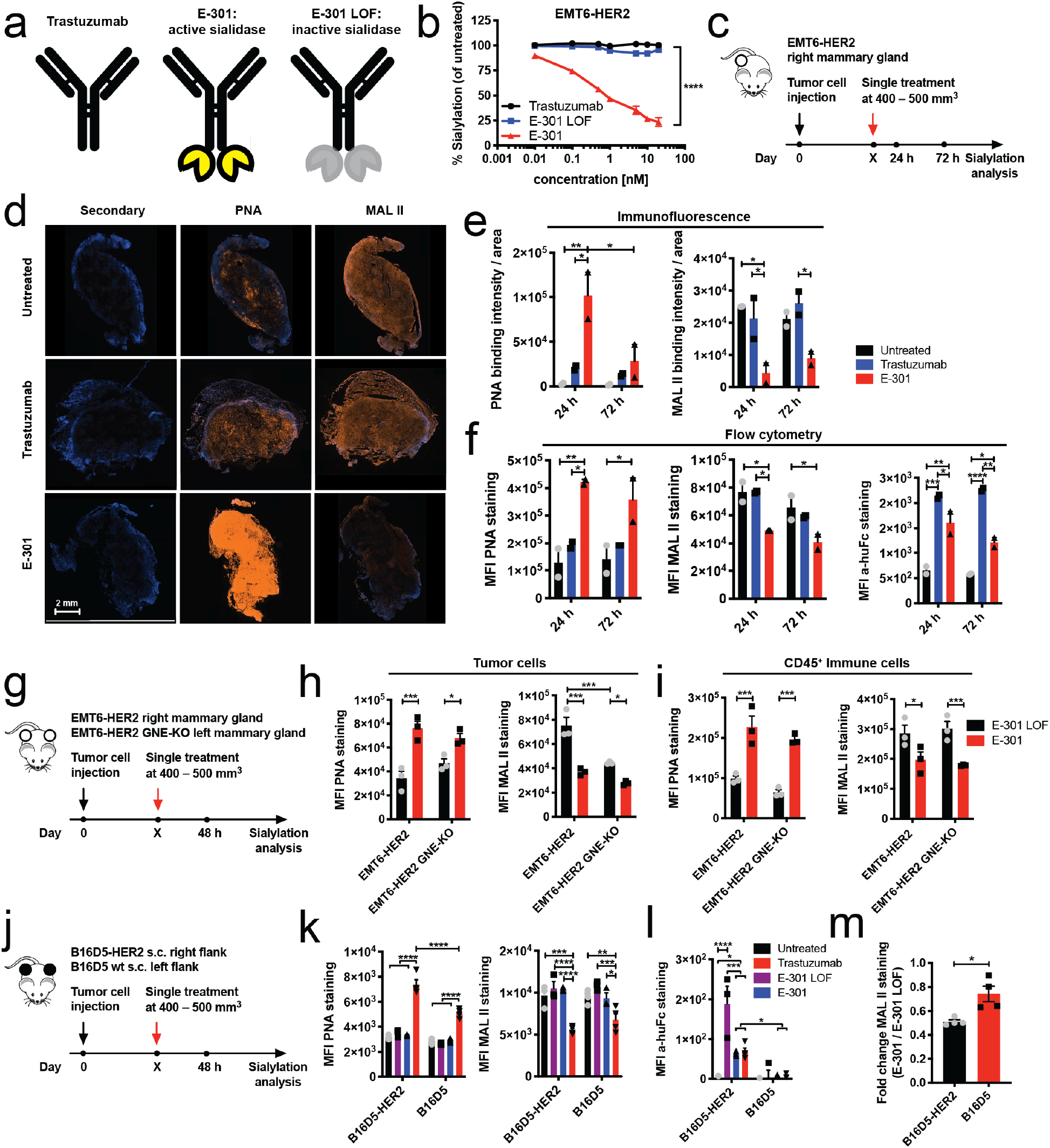
Tumor-targeted sialidase effectively desialylates the tumor microenvironment. **a**, Schematic representation of the tumor-targeted sialidase constructs: trastuzumab, the trastuzumab-sialidase conjugate E-301 and the loss-of-function (LOF) mutated, enzymatically inactive version E-301 LOF. **b**, *In vitro* titration of trastuzumab, E-301 and E-301 LOF on EMT6-HER2 cells; desialylation was assessed by PNA staining 24 h after treatment relative to that after maximal desialylation (n=3). **c**, Experimental setup in order to test *in vivo* desialylation: Mice carrying established (500 mm^3^) intramammary EMT6-HER2 tumors were treated with a single dose of PBS, trastuzumab and E-301 (10 mg/kg) i.p. and desialylation assessed at 24 h and 72 h post treatment. **d**, Representative immunofluorescence images of untreated, trastuzumab- and E-301-treated EMT6-HER2 tumors at 24 h post-treatment, stained with anti-human Fc secondary, PNA and MAL II. **e**, Quantification of immunofluorescence staining with PNA and MAL II; the sum of the staining intensity was normalized to the respective DAPI-stained area. **f**, Analysis of intramammary EMT6-HER2 tumor sialylation (same tumors as in Fig. 2e) by flow cytometry of lectin-stained tumor cell suspensions at 24 h and 72 h after treatment. Geometric mean fluorescence intensities (MFIs) of PNA, MAL II and secondary anti-human Fc staining (d, f, n=2 mice per group). **g**, Experimental setup comparing desialylation of intramammary wildtype and GNE-KO EMT6-HER2 tumors: Mice carrying established (500 mm^3^) intramammary wildtype and GNE-KO EMT6-HER2 tumors were treated with a single dose of E-301 LOF and E-301 (10 mg/kg) i.p. and desialylation assessed at 48 h post treatment. **h**, Geometric MFIs of PNA and MAL II staining of tumor cells. **i**, Geometric MFIs of PNA and MAL II staining of tumor-infiltrating CD45^+^ cells (h, i, n=3 mice per group). **j**, Experimental setup comparing desialylation of subcutaneous B16D5 and B16D5-HER2 tumors at 48 h after i.p. treatment with a single dose of trastuzumab, E-301 or E-301 LOF (10 mg/kg). **k**, Geometric MFIs of PNA, MAL II and **l**, secondary anti-human Fc staining of tumor cells. **m**, Fold change in the geometric MFI of MAL II staining after E-301 treatment relative to E-301 LOF treatment (k–m, n=4). n indicates the number of biological replicates. Error bars represent the mean ± standard error of the mean (s.e.m.). Statistical analyses were performed using two-way ANOVAs with post hoc Sidak’s test an and unpaired two-tailed Student’s *t*-test was used to assess fold change differences in Fig. 2m. * *P* ≤ 0.05, ** *P* ≤ 0.01, *** *P* ≤ 0.001, **** *P* ≤ 0.0001.

Then, we wanted to assess if direct interactions between the sialidase’s glycan-binding domains and tumor sialoglycans might influence sialidase-targeting. To this end, we used EMT6-HER2 cells lacking the rate-limiting enzyme for sialic acid biosynthesis, UDP-GlcNAc 2-epimerase (GNE, GNE-KO) and confirmed comparable expression of the HER2 antigen (Extended Data Fig. 2f). We then injected wildtype EMT6-HER2 and EMT6-HER2 GNE-KO cells into opposing mammary glands of individual mice and again treated mice carrying established tumors with a single dose of E-301 or E-301 LOF (Fig. 2g). At 48 h post-treatment, the wildtype EMT6-HER2 tumors showed strong desialylation, as evidenced by an increase in PNA and a decrease in MAL II staining. The GNE-KO tumors, while already presenting reduced sialylation at baseline, showed a slight further desialylation after treatment (Fig. 2h, Extended Data Fig. 2g), possibly reflecting scavenging of sialic acids form the environment. In contrast, CD45^+^ immune cells showed equally strong loss of sialylation in both wildtype and GNE-KO tumors (Fig. 2h, i). Similar levels of staining with the mannose-binding lectin Concavalin A (ConA) across all groups confirmed the observed changes to be specific to sialic acid-containing glycans (Extended Data Fig. 2h).

We further assessed the capacity of E-301 to target and desialylate HER2-expressing melanoma B16D5 (B16D5-HER2) tumors. C56BL/6 mice were subcutaneously injected with B16D5-HER2 tumor cells in the right and parental B16D5 cells in the left flank (Fig. 2j). Whereas desialylation was detected in both tumor types when treated with E-301 (Fig. 2k), it was significantly more pronounced in the B16D5-HER2 tumors than in the B16D5 tumors (Fig. 2m, Extended Data Fig. 2i). Notably, while the presence of all compounds could be detected in the B16D5-HER2 tumors by staining for human Fc, no staining was observed in the B16D5 tumors (Fig. 2l), suggesting a prolonged retention of the constructs in the HER2-expressing tumors. Again, staining with ConA confirmed sialic acid-specificity of these changes (Extended Data Fig. 2j). These findings demonstrate the feasibility of using E-301 to target systemically administered sialidase into HER2 expressing tumors and confirm effective desialylation of the tumor microenvironment.

### Tumor-targeted sialidase inhibits tumor growth by activating the adaptive immune system

Our next aim was to test the therapeutic efficacy of systemic E-301 treatment in different HER2-expressing tumor models. As a first model, we used the orthotopic, intramammary EMT6-HER2 model (Fig. 3a). Tumor growth was delayed and survival prolonged in mice treated with four doses of E-301, and 1 tumor out of 12 was rejected (Fig. 3b, c). Re-challenge of the tumor-free mouse with either wildtype EMT6 or EMT6-HER2 cells in each flank, both resulted in no tumor growth, indicating the development of immunological memory not restricted to the antigenicity of HER2 (Fig. 3d). The body weights of treated mice increased similarly over the course of the experiment among all treatment groups, suggesting no signs of acute toxicity (Extended Data Fig. 3a). Similarly, B16D5-HER2 tumor growth was clearly delayed and survival prolonged after E-301 monotherapy (Fig. 3e–g). Again, no signs of acute toxicity of the treatment could be observed (Extended Data Fig. 3b). Seeing the development of immunological memory following E-301-induced tumor rejection, we performed an antibody-mediated depletion experiment to test the involvement of the adaptive immune system. We found that depletion of CD8^+^ T cells fully abrogated the delay in tumor growth by E-301 (Fig. 3h, i). These data show that therapeutic tumor-targeted desialylation is efficacious in delaying tumor growth in different mouse models by activating the adaptive immune system and induces the generation of immunological memory.

**Figure 3.**
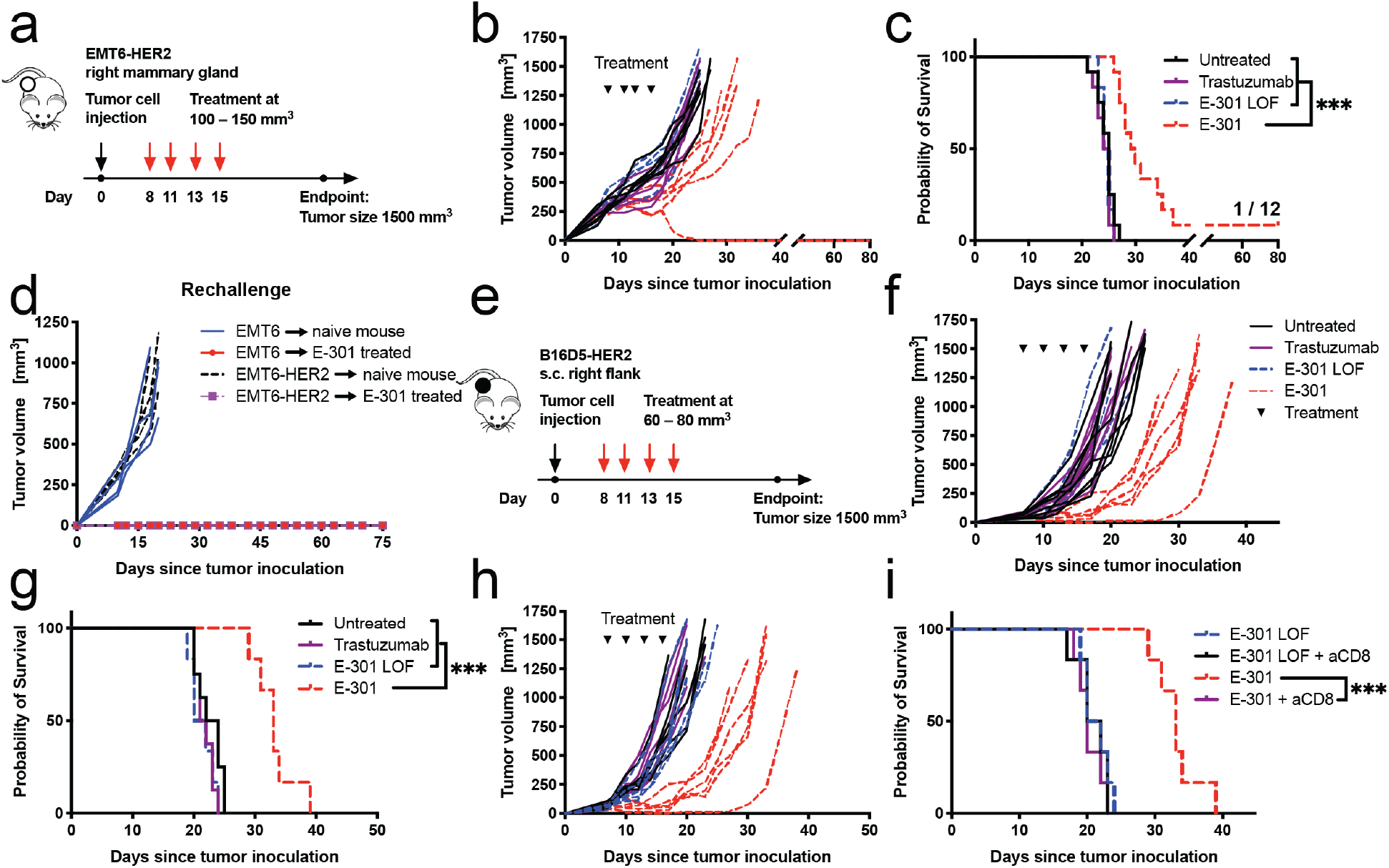
Tumor-targeted sialidase inhibits tumor growth by activating the adaptive immune system. **a**, Experimental setup: Mice carrying intramammary EMT6-HER2 tumors were treated i.p. with four doses of 10 mg/kg trastuzumab, E-301 LOF and E-301, beginning at a tumor size of approximately 100 mm^3^. **b**, Growth of individual intramammary EMT6-HER2 tumors treated with trastuzumab, E-301 LOF or E-301 (n=6–8). **c**, Survival of mice bearing intramammary EMT6-HER2 tumors treated with trastuzumab, E-301 LOF or E-301 (pooled from two experiments, n=12 mice per group). **d**, Rechallenge of the tumor-free mouse from Fig. 3c and tumor naïve control mice with subcutaneous EMT6 or EMT6-HER2 tumor cells in each flank respectively (n=1–4 mice per group). **e**, Experimental setup: Treatment of mice carrying subcutaneous B16D5-HER2 tumors with four doses of trastuzumab, E-301 LOF or E-301 (10 mg/kg) i.p., with the first dose administered once the tumor size reached ∼80 mm^3^. **f**, Growth of individual subcutaneous B16D5-HER2 tumors treated with trastuzumab, E-301 LOF or E-301. **g**, Survival of mice bearing B16D5-HER2 tumors treated with trastuzumab, E-301 LOF or E-301 (f, g, n=6–8 mice per group). **h**, Growth of individual B16D5-HER2 tumors treated with E-301 LOF or E-301 after CD8^+^ T cell depletion. **i**, Impact of CD8^+^ T cell depletion on the survival of mice bearing B16D5-HER2 tumors treated with E-301 LOF or E-301. (h, i, n=6–8 mice per group). n indicates the number of biological replicates. Statistical analyses were performed using the log-rank (Mantel–Cox) test or the Gehan-Wilcoxon test, followed by Bonferroni’s correction for multiple comparisons for all survival analyses. * *P* ≤ 0.05, ** *P* ≤ 0.01, *** *P* ≤ 0.001, **** *P* ≤ 0.0001.

### Tumor-targeted desialylation repolarizes tumor-associated macrophages

In order to dissect the cellular and molecular mechanism by which targeted desialylation augments anti-cancer immune responses, we injected mice carrying palpable subcutaneous B16D5-HER2 tumors i.p. with two doses of either E-301 or E-301 LOF alone or in combination with PD-1 and CTLA-4 blocking antibodies and performed single-cell RNA sequencing (scRNA-seq) on CD45^+^ tumor-infiltrating immune cells 7 days after the first treatment (Fig. 4a). Globally, treatment with E-301 induced distinct changes in populations of both myeloid cells and lymphocytes (Fig. 4b, c, Extended Data Fig. 4a). Most prominently, E-301 affected macrophage populations, resulting in decreased frequencies of clusters 3 and 6 (Fig. 4c, highlighted in blue) and an increase in cluster 14 (Fig. 4c, highlighted in red). Subsetting and reclustering of all macrophages yielded 21 TAM clusters (Fig. 4d), which again showed striking differences following desialylation (Fig. 4e, f). Tumors from untreated and E-301 LOF-treated mice predominantly contained immunosuppressive TAMs expressing high levels of genes characteristic of alternatively activated (M2 polarized) and pro-angiogenic macrophages, such as *Arg1* or *Mrc1* (CD206) (clusters 5, 6 and 11, Fig. 4g, Expanded Data Fig. 4b). In contrast, E-301 treated tumors contained fewer immunosuppressive TAMs and instead displayed a distinct shift towards macrophages predominantly expressing anti-tumoral effector molecules, such as *Il1b, Tnf, Cd80* and *Arg2* (Fig. 4h, clusters 2 and 13, highlighted in red).

**Figure 4.**
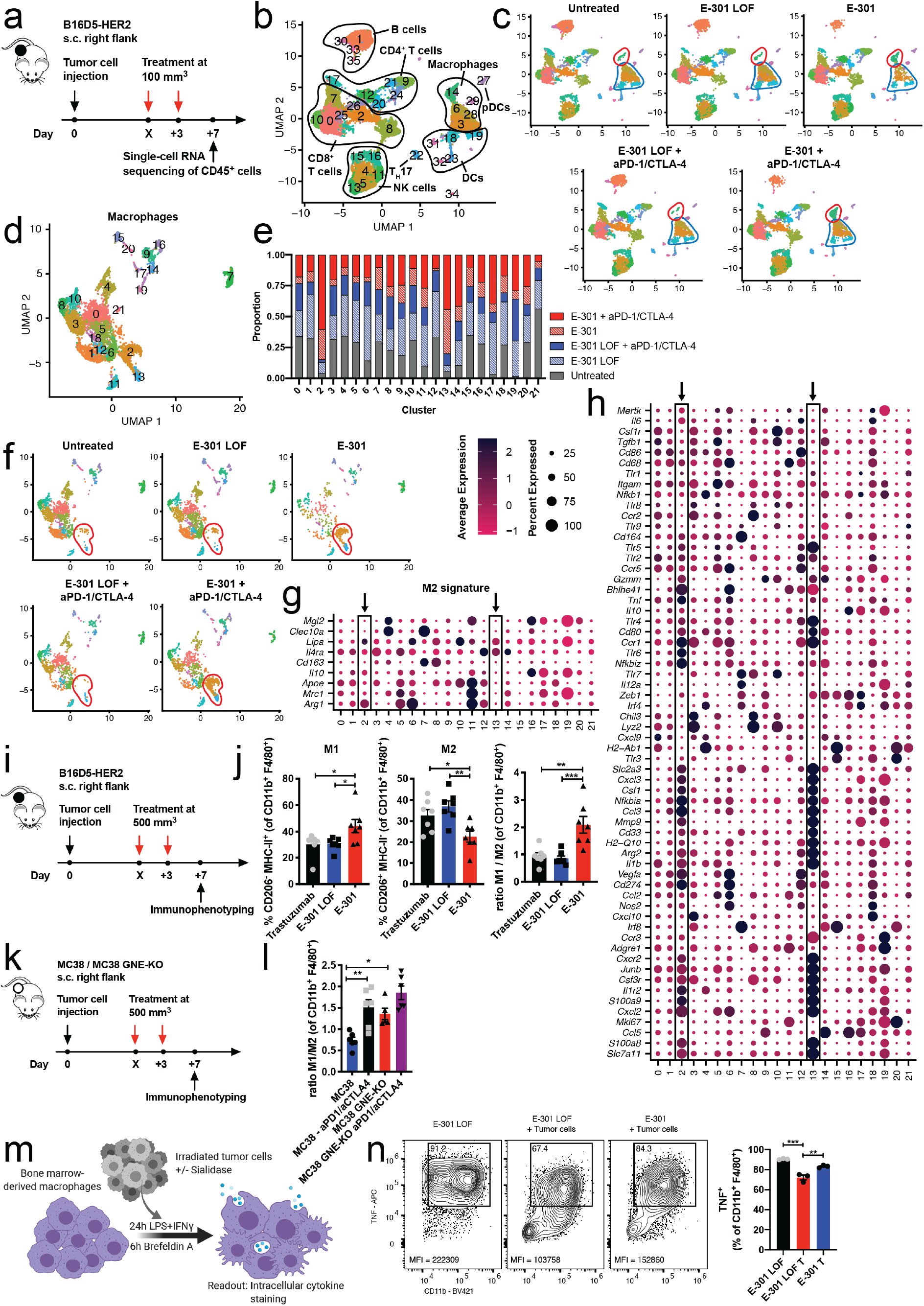
Tumor-targeted desialylation repolarizes tumor-associated macrophages. **a**, Experimental setup for single-cell RNA sequencing (scRNA-seq) of immune infiltrates after E-301 treatment: Mice bearing palpable (100 mm^3^) subcutaneous B16D5-HER2 tumors were treated with two doses of 10 mg/kg E-301 LOF or E-301, alone or in combination with anti-PD-1/CTLA-4 antibodies. CD45^+^ tumor-infiltrating immune cells were isolated and sorted at 7 days after the first injection for scRNA-seq. **b**, scRNA-seq gene expression data was processed, sorted into clusters and is presented in a dimensional reduction projection (UMAP), showing different immune cell populations. Labels have been added based on expression of marker genes. **c**, UMAP projections are shown separated by condition. Clusters 3 and 6 are highlighted in blue, cluster 14 in red. **d**, Subclustering of all macrophages. **e**, UMAP projections of macrophages are shown separated by condition. Clusters 2 and 13 are highlighted in red. **f**, Contribution of each condition to each macrophage cluster. **g**, Dot plot representation of differentially expressed genes between the macrophage clusters, genes characteristic for M2 polarization, or in **h**, reflecting more general macrophage function. Size reflects the percentage of each cluster expressing a given gene, average scaled expression is indicated on the color gradient. Clusters 2 and 13 are boxed in and highlighted with arrows (a–h, n=5 pooled mice per condition). **i**, Experimental setup for immunophenotyping of changes in immune infiltrates after E-301 treatment: Mice bearing established (500 mm^3^) subcutaneous B16D5-HER2 tumors were treated with two doses of 10 mg/kg trastuzumab, E-301 LOF or E-301 and immune infiltrates analyzed after 7 days by flow cytometry. **j**, Frequencies of CD206^-^MHC-II^+^ (M1) and CD206^+^MHC-II^-^ (M2) cells among CD11b^+^F4/80^+^ tumor-associated macrophages. Ratio of M1 to M2 macrophages in CD11b^+^F4/80^+^ cells (n=7). **k**, Experimental design: Mice carrying established (approx. 500 mm^3^) subcutaneous wildtype and GNE-KO MC38 tumors were treated i.p. with two doses of 10 mg/kg anti-PD-1 and anti-CTLA-4 antibodies. 7 days after the first treatment, tumors were resected and immunophenotyped. Same tumors as in Fig. 1k. **l**, Ratio of M1 to M2 macrophages among CD11b^+^F4/80^+^ cells (n=5–6). **m**, Experimental setup for *in vitro* coculture of bone marrow-derived macrophages (BMDMs), irradiated B16D5-HER2 tumor cells and sialidase. **n**, Representative flow cytometry dot plots of anti-TNF and anti-CD11b staining. Gates show percentage of TNF^+^ cells, MFI indicates the mean fluorescence intensity of TNF staining in the TNF^+^ population. Quantification of TNF^+^ cells among CD11b^+^F4/80^+^ BMDMs (n=3). n indicates the number of biological replicates. Error bars represent the mean ± standard error of the mean (s.e.m.). Statistical analyses were performed using one-way ANOVA with post hoc Tukey’s test. * *P* ≤ 0.05, ** *P* ≤ 0.01, *** *P* ≤ 0.001, **** *P* ≤ 0.0001.

Treatment with E-301 also affected intratumoral NK and T cell populations (Expanded Data Fig. 4c, f). Among NK cells, desialylation increased the frequencies of all anti-tumoral NK cell subsets (Extended Data Fig. 4d, e). Similarly, analysis of all T cells (Extended Data Fig. 4f), revealed decreases in naïve CD4^+^ T cells (CD4^+^ T_N_, cluster 7) and exhausted CD8^+^ T cells (CD8^+^ T_EX_, cluster 8), and the expansion of effector CD8^+^ T cells (CD8^+^ T_EFF_, clusters 5 and 13) and both T_h_1 and T_h_2 CD4^+^ T cells (CD4^+^ T_h_1, T_h_2, clusters 4 and 6; Extended Data Fig. 4g, h).

We confirmed our findings by flow cytometric immunophenotyping of established B16D5-HER2 tumors, treated with two doses of E-301, E-301 LOF or trastuzumab (Fig. 4i). We found a significantly higher absolute number of both total CD45^+^ cells and CD8^+^ T cells in tumors treated with E-301 compared with the control-treated tumors (Extended Data Fig. 4i). In agreement with our scRNA-seq results, we further observed an increase in M1 polarized and a decrease in M2 polarized TAMs, evidenced both by MHC-II, CD206 staining (Fig. 4j) and by the expression of the activation marker CD80 (Extended Data Fig. 4j). Consequently, the M1/M2 ratio was increased upon E-301 treatment (Fig. 4j). Similarly, DCs displayed an increase in the activation marker CD40 (Extended Data Fig. 4k). Finally, CD8^+^ T cells showed increased expression of the effector molecules granzyme B and Ki67 (Extended Data Fig. 4l). Similar findings were made in EMT6-HER2 tumors (Extended Data Fig. 4m), with E-301-treatment resulting in a shift from M2 to M1 polarized TAMs (Extended Data Fig. 4n) and an increase in granzyme B and Ki67 expressing CD8^+^ T cells (Extended Data Fig. 4o). Analysis of macrophages from ICB-treated wildtype and GNE-KO tumors (Fig. 4k) revealed a similar repolarization of TAMs in the desialylated tumors, which was further increased by ICB (Fig. 4l). Interestingly, restimulation of NK cells in single-cell suspensions of those tumors showed an increase in the frequency of IFNγ^+^ NK cells in the GNE-KO tumors compared to the wildtype tumors, but no further increase in combination with ICB (Extended Data Fig. 4p). Multiplex analysis of serum cytokine levels from E-301 treated mice carrying B16D5-HER2 tumors revealed significantly higher levels of cytokines including IL-13, IL-3, IFNa, IL-6 and IL-27 (Extended Data Fig. 4q). Together, these results demonstrate profound remodeling of the tumor immune microenvironment after therapeutic desialylation by E-301. The specific increase of anti-tumoral macrophage populations results in an overall shift in intratumoral macrophage polarization and augments the generation of an antitumoral CD8^+^ T cell response.

In order to model our observation of repolarization of TAMs after *in vivo* tumor desialylation, we set up an *in vitro* co-culture system of bone-marrow derived macrophages (BMDMs) with irradiated B16D5-HER2 tumor cells (Fig. 4m). We found tumor cells to inhibit pro-inflammatory activation of BMDMs, measured by intracellular TNF staining, which could be partly rescued by E-301 treatment, but not by E-301 LOF (Fig. 5n). Similarly, phagocytosis, measured by the macrophage-intrinsic fluorescence intensity of CypHer5E, a pH-sensitive fluorophore, was increased after i.p. injection of desialylated CypHer5E-labelled MC38 GNE-KO tumor cells, compared to labelled wildtype MC38 cells (Extended Data Fig. 4r). This goes in line with previous work and the description of sialic acids as “don’t eat me” signals^12^ and supports our findings linking tumor sialylation to a pro-tumorigenic TAM polarization.

**Figure 5.**
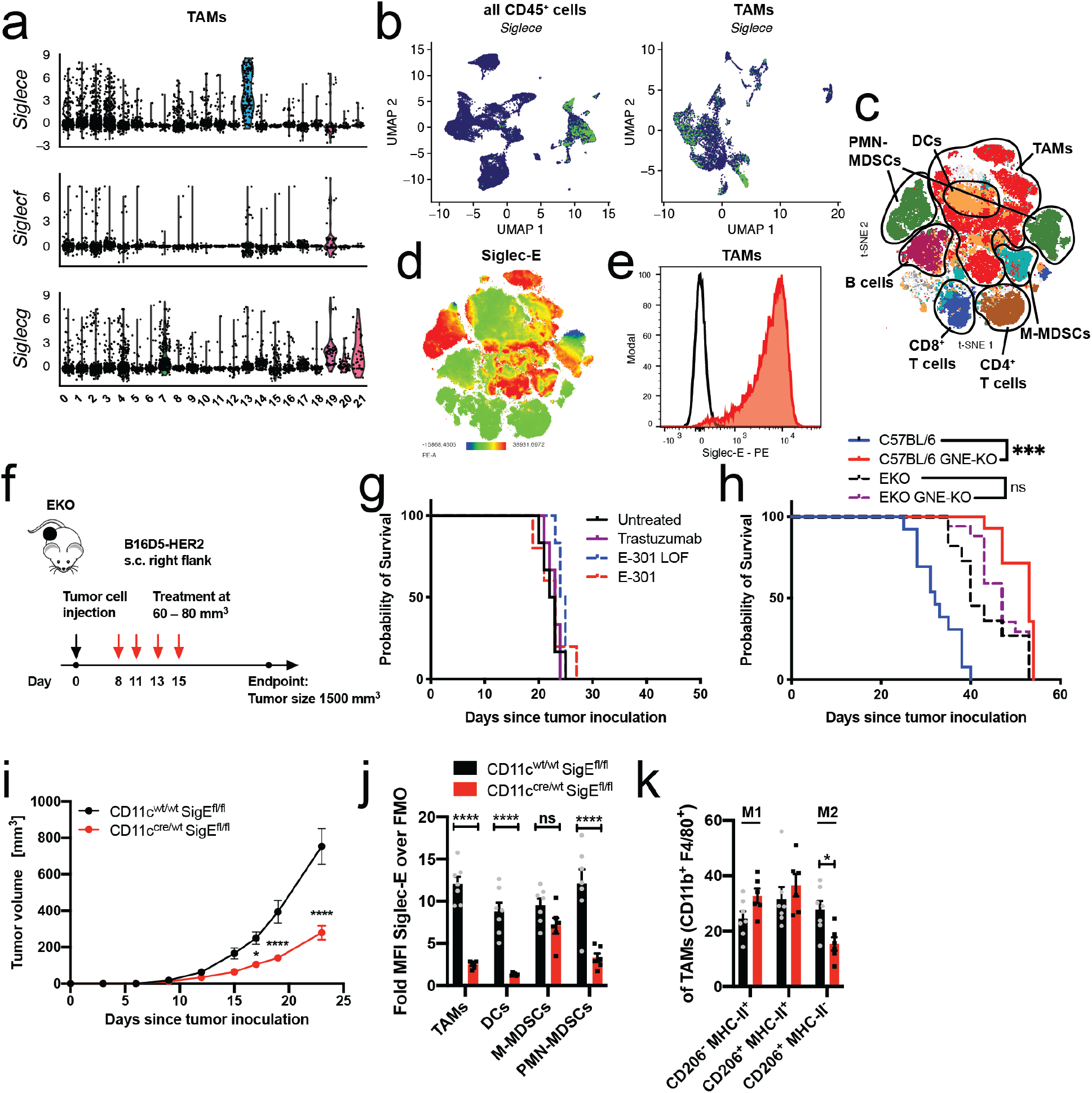
Efficacy of tumor-targeted sialidase is dependent of Siglec-E on TAMs. **a**, Expression of *Siglece, Siglecf* and *Siglecg* in all macrophage clusters from scRNA-seq data (Fig. 4a). Sample distribution shown as violin plots. **b**, UMAP projections of all CD45^+^ cells and all macrophages from scRNA-seq data. Expression of *Siglece* is shown as a color gradient from blue (low) to green (high). **c**, t-distributed stochastic neighbor embedding (t-SNE) projection of multicolor flow cytometric immunophenotyping of B16D5-HER2 and EMT6-HER2 tumors. Cell populations have been assigned based on maker expression (Extended Data Fig. 5a, b). **d**, Staining intensity for Siglec-E is shown as a color gradient from blue (low) to red (high). **e**, Representative histogram of Siglec-E staining on CD11b^+^F4/80^+^ tumor-associated macrophages from B16D5-HER2 tumor. Isotype control staining is shown as empty histogram, anti-Siglec-E staining in red. **f**, Experimental setup: Mice deficient in Siglec-E (EKO) bearing subcutaneous B16D5-HER2 tumors were treated i.p. with four doses of trastuzumab, E-301 LOF or E-301 (10 mg/kg) beginning once the tumor size reached ∼80 mm^3^. **g**, Survival of EKO mice bearing subcutaneous B16D5-HER2 tumors after trastuzumab, E-301 LOF or E-301 treatment (n=6 mice per group). **h**, Survival of wildtype C57BL/6 and EKO mice after subcutaneous injection of MC38 wildtype and GNE-KO tumor cells (n=13–17 mice per group). **i**, Tumor growth of subcutaneously injected MC38 cells in Siglec-E^fl/fl^ (Elox) mice crossed to CD11c-Cre mice. CD11c^cre/wt^Siglec-E^fl/fl^ mice, lacking Siglec-E in all CD11c-expressing cells are compared to their wildtype CD11c^wt/wt^Siglec-E^fl/fl^ littermate controls (n=7–8). **j**, Siglec-E expression on different tumor-infiltrating myeloid immune cells in CD11c-Cre Elox mice compared to littermate control mice, by flow cytometry. Siglec-E expression shown as fold change over control staining. **k**, Frequencies of CD206^-^MHC-II^+^ (M1), CD206^+^MHC-II^+^ and CD206^+^MHC-II^-^ (M2) macrophages among CD11b^+^F4/80^+^ cells. CD11c^cre/wt^Siglec-E^fl/fl^ mice, lacking Siglec-E in all CD11c-expressing cells are compared to their wildtype CD11c^wt/wt^Siglec-E^fl/fl^ littermate controls (m, n, n=7–8). n indicates the number of biological replicates. Error bars represent the mean ± standard error of the mean (s.e.m.). Statistical analyses were performed using the log-rank (Mantel–Cox) test or the Gehan-Wilcoxon test, followed by Bonferroni’s correction for multiple comparisons for all survival analyses. Differences in Fig. 5i–k were tested using two-way ANOVAs with post hoc Tukey’s test. * *P* ≤ 0.05, ** *P* ≤ 0.01, *** *P* ≤ 0.001, **** *P* ≤ 0.0001.

### Efficacy of tumor-targeted sialidase is dependent of Siglec-E on TAMs

Next, we set out to identify the mechanism by which desialylation affects the polarization of intratumoral macrophages. To that end, we analyzed and compared the expression of the most common inhibitory CD33-related Siglecs, *Siglece, Siglecf* and *Siglecg* in all TAMs in our scRNA-seq data and found *Siglece* to be the most prominently expressed (Fig. 5a). Besides being broadly expressed on most TAMs, *Siglece* was most highly expressed on the anti-tumorigenic TAM cluster 13, which was strongly increased in E-301-treated tumors. This falls in line with previous results showing it to be the most broadly expressed inhibitory Siglec in mice^5,18^. Further comparison of *Siglece* expression among all CD45^+^ cells confirmed it to be predominantly expressed on macrophages (Fig. 5b). To validate this finding on protein level and in order to include granulocytes, which are commonly underrepresented in scRNA-seq, we used multicolor immunophenotyping of single-cell suspensions from both B16D5-HER2 and EMT6-HER2 tumors. T-stochastic next neighbor (t-SNE) dimensional reduction of 10 concatenated samples, allowed the identification of all major intratumoral immune cell types, including M-MDSCs and PMN-MDSCs (Fig. 5c, Extended Data Fig. 5c, d). Overlay of the intensity of anti-Siglec-E staining (Fig. 5d, Extended Data Fig. 5e), revealed the strongest expression of Siglec-E on TAMs and PMN-MDSCs (Fig. 5e). Among CD11b^+^ cells, Siglec-E expression coincided with the strongest expression of F4/80 and notably CD11c expression (Extended Data Fig. 5b).

To validate the role of Siglec-E as the receptor for tumor sialylation, we assessed the growth of subcutaneous B16D5-HER2 tumors in C57BL/6 mice lacking Siglec-E (EKO, Fig. 5f). Treatment of EKO mice carrying B16D5-HER2 tumors with E-301, E-301 LOF or trastuzumab did not delay tumor growth (Fig. 5g, Extended Data Fig. 5f), supporting the role of Siglec-E in mediating the effects of desialylation. This finding was in accordance with earlier experiments^14^. To further corroborate this finding, we used MC38 GNE-KO tumor cells as a genetic model of desialylation. Again, while growth of MC38 GNE-KO tumors was significantly delayed and survival prolonged in wildtype C57BL/6 mice, when compared to parental MC38 tumors, no difference in tumor growth between MC38 and desialylated MC38 GNE-KO tumors was observed in EKO mice (Fig. 5h, Extended Data Fig. 5g).

In a next step, we wanted to test the effect TAM-specific loss of Siglec-E expression. To this end, we generated a new conditional mouse knockout strain, by generating mice carrying floxed Siglec-E alleles (Siglec-E^fl/fl^, Elox). Crossing Elox mice with CD11c-Cre mice, we obtained a conditional knockout of Siglec-E on all CD11c^+^ cells. While CD11c is widely used as a marker for DCs, it is also described as a marker specific for tumor-associated macrophages^19^, which we were able to confirm by finding the highest expression of Siglec-E among TAMs to coincide with the highest expression of CD11c. (Fig. 5c, Extended Data Fig. 5b). Growth of MC38 tumors was delayed in Siglec-E^fl/fl^xCD11c^Cre/wt^ mice, lacking Siglec-E on CD11c^+^ cells, compared to their Siglec-E^fl/fl^xCD11c^wt/wt^ control littermates (Fig. 5i). Flow cytometric immunophenotyping of Siglec-E expression on tumor-infiltrating immune cells confirmed loss of Siglec-E on both DCs and TAMs, as well as on PMN-MDSCs (Fig. 5j, Extended Data Fig. 5h), while no difference was detected on lymphoid cells (Extended Data Fig. 5i). Notably, while intratumoral macrophages and DCs showed comparable expression of Siglec-E, splenic macrophages expressed much less Siglec-E than splenic DCs and showed no reduction in Siglec-E expression in Siglec-E^fl/fl^xCD11c^Cre/wt^ mice, while splenic DCs remained affected, further supporting the use of CD11c-Cre as a driver for tumor-specific macrophage knockout (Extended Data Fig. 5j). In order to address potential contribution of DCs to the tumor growth delay in the CD11c-Cre model, we crossed Elox mice with XCR1-Cre mice to obtain a cDC1-specific deletion of Siglec-E. In contrast to the delay observed in the CD11c-specific knockout mice, XCR1-Cre mice showed no difference in tumor growth between Siglec-E^fl/fl^xXCR1^Cre/wt^ mice and Siglec-E^fl/fl^xXCR1^wt/wt^ littermate controls (Extended Data Fig. 5k). Taken together, these experiments identify Siglec-E on TAMs as the receptor for tumor sialylation. Deletion of Siglec-E abrogated the effects of both genetic and therapeutic desialylation and targeting of Siglec-E on CD11c^+^ TAMs delayed tumor growth and enhanced antitumor immunity.

### Targeting tumor sialylation or Siglec-E synergizes with immune checkpoint blockade

We then asked if tumor-targeted desialylation could be combined with PD-1 and CTLA-4 blockade in the B16D5-HER2 model (Fig. 6a). Combination of E-301 with PD-1- and CTLA-4-blocking antibodies augmented inhibition of tumor growth and delayed time to progression compared with both E-301 or ICB in combination with E-301 LOF (Fig. 6b, c). This finding corroborated the synergism between desialylation and ICB which we had observed using several genetic models of desialylation (Fig. 1h–j). Together, these data clearly demonstrate a synergistic effect of targeting tumor sialylation and PD-1/CTLA-4 blockade.

**Figure 6.**
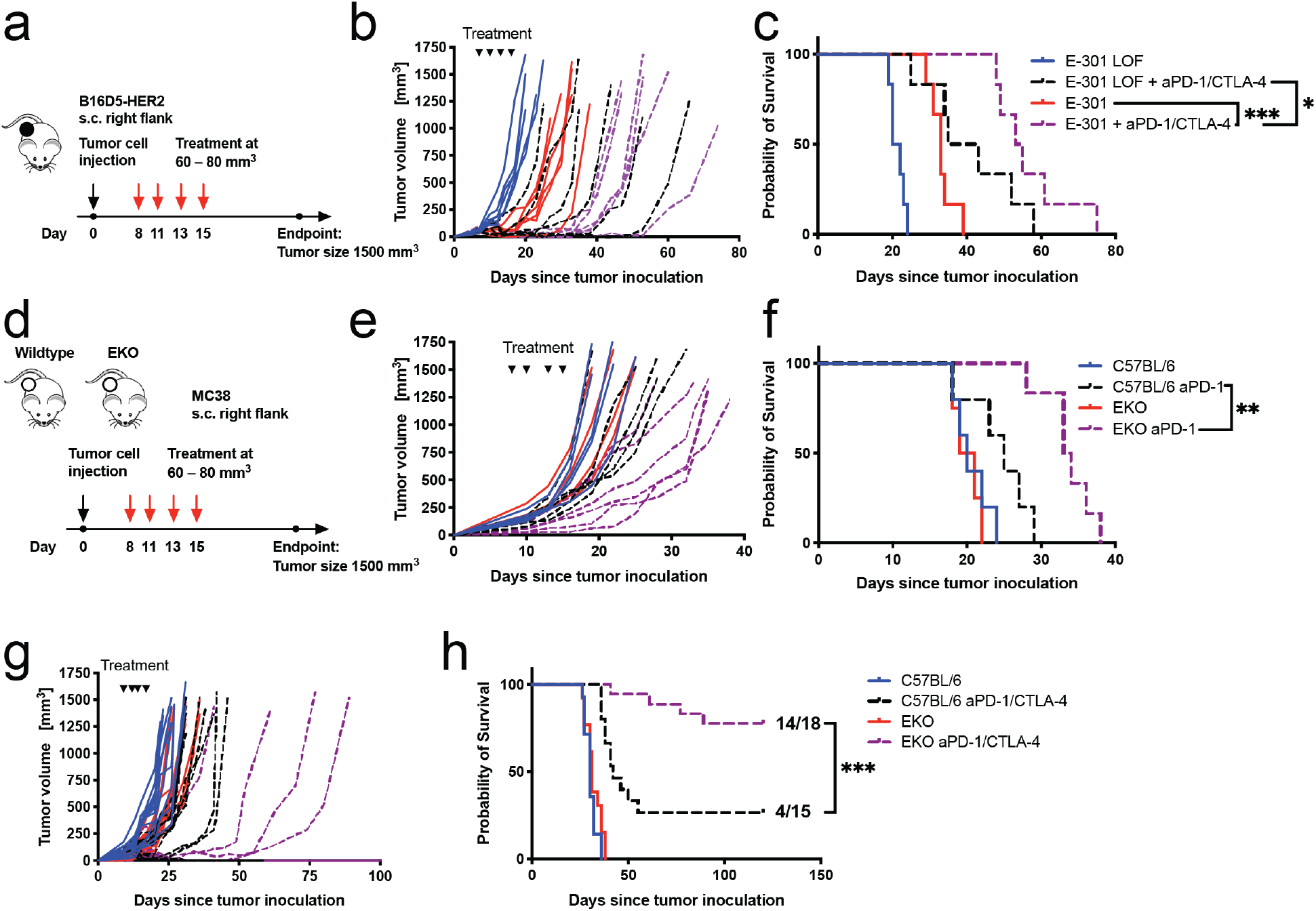
Targeting tumor sialylation or Siglec-E synergizes with ICB. **a**, Experimental setup: Treatment of mice carrying subcutaneous B16D5-HER2 tumors with four doses of trastuzumab, E-301 LOF or E-301 (10 mg/kg) i.p., with the first dose administered once the tumor size reached ∼80 mm^3^. **b**, Growth of individual B16D5-HER2 tumors treated with E-301 LOF or E-301 in combination with anti-PD-1/CTLA-4 antibodies. **c**, Survival of mice bearing B16D5-HER2 tumors treated with E-301 LOF or E-301 in combination with anti-PD-1/CTLA-4 antibodies (b, c, n=6–8 mice per group). **d**, Experimental setup for the treatment of wildtype C57BL/6 and EKO mice bearing subcutaneous MC38 tumors with four doses of anti-PD-1 or/and anti-CTLA-4 antibodies (10 mg/kg) i.p. beginning once the tumor size reached ∼80 mm^3^. **e**, Effect of anti-PD-1 ICB on the growth of individual MC38 tumors in wildtype C57BL/6 and EKO mice. **f**, Survival of wildtype C57BL/6 and EKO mice bearing MC38 tumors after treatment with PD-1 blockade (e, f, n=4–6 mice per group). **g**, Effect of anti-PD-1 and anti-CTLA-4 ICB on the growth of individual MC38 tumors in wildtype C57BL/6 and EKO mice. **h**, Survival of wildtype C57BL/6 and EKO mice bearing MC38 tumors after treatment with PD-1 and CTLA-4 blockade (g, h, n=13– 18 mice per group). n indicates the number of biological replicates. Statistical analyses were performed using the Gehan-Wilcoxon test followed by Bonferroni’s correction for multiple comparisons for all survival analyses. * *P* ≤ 0.05, ** *P* ≤ 0.01, *** *P* ≤ 0.001, **** *P* ≤ 0.0001.

Based on our finding that Siglec-E mediates the effect of therapeutic desialylation, we wanted to test if loss of Siglec-E would similarly synergize with ICB. To that end, we inoculated MC38 tumors in both wildtype C57BL/6 and EKO mice (Fig. 6d). PD-1 blockade was significantly more efficacious in delaying tumor growth and prolonging survival in EKO mice than in wildtype C57BL/6 mice (Fig. 6e, f). This effect was further enhanced when a combination of PD-1- and CTLA-4-blockade was used, with 14 of 18 (78%) EKO mice rejecting their tumors, compared to 4 of 15 (27%) wildtype mice (Fig. 6g, h). These findings further support the sialoglycan-Siglec axis as a target for immunotherapy, as both genetic and therapeutic desialylation, as well as loss of Siglec-E synergize with ICB.

## Discussion

Tumor sialylation dampens anti-tumor immune responses and targeting the sialoglycan–Siglec axis has been proposed as a new immunotherapeutic approach. Our data provides mechanistic evidence for tumor sialylation-mediated immune suppression, demonstrates the feasibility of therapeutic desialylation and its potential for use in combination with classical ICB. We show that inhibitory Siglec-E on TAMs is the main receptor for cancer-associated sialic acids and demonstrate specific anti-tumoral repolarization of TAMs upon tumor desialylation or loss of Siglec-E (Graphical abstract, Extended Data Fig. 6).

We recently described the upregulation of human Siglec-9 and murine Siglec-E on tumor-infiltrating T cells in humans and mice^6^. This upregulated expression dampens T cell activation in the presence of Siglec-ligands, which can be reversed by blockade of sialoglycan-binding. It is conceivable that direct reinvigoration of T cell effector function in the tumor microenvironment might contribute to the results obtained in this work. However, we here identified Siglec-E expression on TAMs as the dominant factor mediating immune suppression due to hypersialylation. Specifically, we demonstrate that loss of Siglec-E abrogated the efficacy of therapeutic and genetic desialylation and that knockout of Siglec-E on TAMs was sufficient to delay tumor growth and repolarize macrophages. We observe enhanced phagocytosis of desialylated tumor cells by macrophages, in line with the recent description of human Siglec-10 as a receptor for ‘don’t eat me’ signals in cancer^12^. Other work using desialylated tumor cell lines or intratumoral administration of a fluorinated sialic acid mimetic, blocking *de novo* biosynthesis of sialoglycans, revealed similar anti-tumor immunity promoting effects^6,20,21^. The uptake of sialylated antigens by DCs was shown to enhance the induction of Tregs through engagement of Siglec-E, which might reflect the interactions between DCs and naïve CD4^+^ T cells in the hypersialylated tumor microenvironment^22^. Notably, we observe a significant reduction in the frequency of naïve CD4^+^ cells upon E-301 treatment (Expended Data Fig. 4f). Besides binding to Siglecs, sialic acid-containing glycans might also inhibit adaptive immunity by acting as alternative ligands for CD28 and inhibiting CD28-CD80 interactions^23^. Additionally, a recent screen for surface proteins inhibiting T cell activation led to the identification of Siglec-15 as a potential target for cancer immunotherapy^24^. Together with our data, these findings suggest a clear role for tumor sialylation in dampening adaptive immune responses, both directly through action on T cells and indirectly by affecting the phenotypes of other tumor-infiltrating immune cell types, such as DCs and TAMs.

Therapeutic targeting of the sialoglycan–Siglec axis can be achieved by either targeting of the sialic acid-containing ligands or the Siglec-receptors. While the later can be accomplished using blocking antibodies, functional redundancy and potential compensation among the human Siglec-repertoire might ultimately dampen the efficacy of such an approach. Therefore, direct targeting of tumor sialylation might prove a more effective option^14^. Blocking tumor hypersialylation with chemical inhibitors of sialic acid biosynthesis results in pronounced desialylation, increased T cell infiltration and CD8^+^ T cell-dependent delay in tumor growth^21^. However, systemic application in mice leads to strong desialylation of all tissues, lasting up to several weeks, and is ultimately fatal, limiting the applicability of this approach to localized intratumoral applications^25^. In addition, complete abrogation of tumor sialylation has been shown to induce apoptosis in CD8^+^ T cells, which might limit the efficacy of such approaches^26^. In contrast, mice systemically injected with *Vibrio cholerae* sialidase only show transient toxicity, reflecting the short-lived nature of enzymatic desialylation^27^. We demonstrate further reduced toxicity by targeting sialidase to tumor-antigens, thereby restricting the sialidase activity to the tumor microenvironment. We suggest this approach to be favorable both in terms of anti-tumor efficacy and potential toxicity.

A limitation of our study is the requirement of a tumor-antigen as the target for therapeutic desialylation. Here, we used HER2 as a well-studied cancer-associated antigen. Although HER2-targeting is effective and has significantly improved the prognosis of patients with HER2 positive breast and gastric cancer^28^, many cancer types lack a selectively and consistently expressed target. Additional work will be needed to specify the requirements for cancer-associated targets for therapeutic desialylation. Of note, the relative contributions of the desialylation of tumor cells and immune cells to the observed anti-tumor activity remain undefined. It remains unclear if the desialylation of tumor cells is an essential prerequisite for the effectiveness of therapeutic desialylation or if targeting of intratumoral immune cells might elicit comparable effects. This could open the possibility of constructing antibody-sialidase constructs against antigens expressed on intratumoral immune cells, expanding the range of potential cancer types.

Our work demonstrates for the first time that therapeutic targeting of tumor sialylation is effective *in vivo* and synergizes with PD-1 and CTLA-4 blockade. Using scRNA-seq, we show mechanistically how therapeutic desialylation repolarizes TAMs towards an anti-tumorigenic phenotype and augments the adaptive anti-tumor immune response. We identify inhibitory Siglec-E on TAMs as the main target of desialylation and provide a strong rationale for the further clinical development of sialoglycan–Siglec targeting agents and their combination with PD-1 and CTLA-4 blocking immunotherapies.

## Materials and Methods

### Mice

All mouse experiments were approved by the local ethics committee (Approval 2747, Basel Stadt, Switzerland) and performed in accordance with the Swiss federal regulations. C57BL/6 and BALB/c mice were obtained from Janvier Labs (France) and bred in-house at the Department of Biomedicine, University Hospital Basel, Switzerland. The Siglec-E-deficient mouse strain (EKO) was received from Dr. Varki (UCSD, San Diego, USA) and had been previously described^29^. EKO mice were bred in-house and backcrossed to the local C57BL/6 strain in heterozygous pairings for more than nine generations. All animals were housed under specific pathogen-free conditions.

### Cells and cell culture

All cell lines were maintained in Dulbecco’s Modified Eagle Medium (DMEM), supplemented with 10% heat-inactivated fetal bovine serum (FBS, PAA Laboratories, Germany), 2 mM L-glutamine, 1 mM sodium pyruvate, 100 µg/mL streptomycin and 100 U/mL penicillin (Gibco, USA). The B16D5-HER2, EMT6-HER2, EMT6-HER2 GNE-KO and MC38 GNE-KO cell lines have been described previously^6,30^. The parental B16F10 cell line was obtained from ATCC (USA) and the generation of B16F10 GNE-KO cells is described below. All cells were cultured at 37°C under 5% CO_2_ atmosphere and cultured for a minimum of 3 passages before being used.

### Tumor models

For tumor growth experiments, 7–11-week-old mice were used. Sex-matched wildtype littermates were used as controls for all experiments involving transgenic mice. Tumor cell injections were performed as described previously^30^. For the subcutaneous MC38 and B16 models (wildtype, GNE-KO, and HER2 expressing), mice were subcutaneously injected with 500,000 tumor cells in phosphate-buffered saline (PBS) into the right flank, or in some cases into both the right and left flank. For the orthotopic EMT6 model, mice were injected with 1,000,000 EMT6 cells in PBS (wildtype, GNE-KO and HER2 expressing) into the right or left mammary gland of female BALB/c mice. Tumor size and overall health score, as well as body weights in some experiments, were measured and monitored three times per week. Perpendicular tumor diameters were measured using a caliper, and tumor volume was calculated according to the following formula: tumor volume (mm^3^)=(d^2^*D)/2, where d and D are the shortest and longest diameters (in mm) of the tumor, respectively. Mice were sacrificed before the size of their tumors reached 1500 mm^3^. Animals that developed ulcerated tumors were sacrificed and excluded from further analysis.

### Treatments

Antibodies for *in vivo* depletion of CD8^+^ T cells – anti-mouse CD8a (53-6.7) – and *in vivo* ICI treatment – anti-mouse PD-1 (RMP1-14) and anti-mouse CTLA-4 (9D9) – were purchased from Bio X Cell (USA). For T cell depletion, anti-CD8 depleting antibody was administered i.p. at 10 mg/kg on days −2, 0, and every 7 days ongoing, relative to the time of tumor cell injection. For efficacy studies, treatments were administered i.p. at 10 mg/kg once the tumors reached an average size of 80–100 mm^3^ and a total of four doses was given every second to third day, unless specified otherwise. For 7-day treatments, tumors were allowed to grow until they reached a size of approximately 500 mm^3^ and treated i.p. at 10 mg/kg with a total of two doses.

### Biologics

The DNA encoding for E-301 and E-301-LOF were cloned into the mammalian expression vector pCEP4 vector (Thermo FisherScientific). HEK 293 cells were then transiently transfected with the DNA constructs using Expi293 Expression system and standard protocols according to the manufacturer’s instructions (Thermo FisherScientific). Purification of E-301 and E-301-LOF was performed directly from transfection harvests using a HiTrap Protein A affinity column (GE Healthcare) and eluted with 1M arginine pH 3.9. Anion-exchange chromatography was used as a secondary purification method and the final product was dialyzed into 1X PBS 7.4. The biochemical characterization of E-301 and E-301-LOF, including purity, Her2 binding affinity, and enzymatic activity, was conducted as described previously (WO2019136167).

### Generation of B16F10 GNE-KO tumor cells

Knockout of *Gne* in B16F10 tumor cells was performed using CRISPR/Cas9 mediated gene editing. Guide RNAs were designed online using E-CRISP (e-crisp.org), synthesized by Microsynth (Switzerland) and cloned into the pSpCas9(BB)-2A-GFP (PX458) vector, a gift from Feng Zhang^31^ (Addgene plasmid #48138; http://n2t.net/addgene:48138; RRID: Addgene_48138, USA). The paternal cell line was transiently transfected and single GFP^+^ cells were sorted into 96-well plates. After their recovery and expansion, individual clones were screened for cell surface sialylation by lectin staining and cells showing reduced staining intensities confirmed to be GNE-KO. Multiple GNE-KO clones were selected and pooled to avoid clonal selection. The wildtype parental cell line, as well as transfected and sorted clones showing normal cell surface sialylation were used as controls. GNE-KO cells were compared to the parental cell line and control clones with regard to their *in vitro* viability and proliferation, as well as their *in vivo* tumor growth, both in immunodeficient NSG mice and wildtype C57BL/6 mice.

### Tumor digests

For the preparation of single cell suspensions from both human and mouse tumors, tumors were collected, surgical specimens were mechanically dissociated and subsequently digested using accutase (PAA Laboratories, Germany), collagenase IV (Worthington, USA), hyaluronidase (Sigma, USA) and DNase type IV (Sigma, USA) for 1 h at 37°C under constant agitation. Cell suspensions were filtered through a 70 µm mesh, and, for the analysis of tumor-infiltrating immune cells, CD45^+^ cells were further enriched by Histopaque-1119 density gradient centrifugation (Sigma, USA). Splenocytes were isolated by mechanical disruption using the end of a 1 mL syringe, filtration through a 70 µm mesh and lysis of red blood cells using RBC lysis buffer (eBioscience, USA). Samples were frozen (in 90% FBS, 10% DMSO) and stored in liquid nitrogen until the time of analysis.

### Multicolor flow cytometry

Flow cytometry was performed on single cell suspensions of cell lines, blood samples, splenocytes and tumor digests. In order to prevent unspecific staining, cells were initially blocked with anti-mouse Fcγ III/II receptor (CD16/CD32) antibodies and dead cells excluded by staining with a fixable live/dead cell-exclusion dye (BioLegend, USA). Then, cell suspensions were stained for cell surface antigens with primary fluorophore-conjugated antibodies for 20 min at 4°C in FACS buffer (PBS, 2% FBS, 0.5 mM EDTA). Stained samples were fixed with IC fixation buffer (eBioscience, USA) until time of analysis. For intracellular antigens (IL-2, IFNγ, TNF, FoxP3 and Ki67), cells were first stained against cell surface antigens, fixed and permeabilized (eBioscience, USA) followed by staining with antibodies directed against intracellular antigens. For intracellular cytokine staining, single cell suspensions of tumor digests were restimulated *ex vivo* with phorbol 12-myristate 13-acetate (PMA, 20 ng/mL) and ionomycin (500 µg/mL) for 6 h, and Brefeldin A (BD Pharmingen, USA) added for the last 4 h. All samples were acquired on a LSR II Fortessa flow cytometer (BD Biosciences, USA) or Cytek Aurora (Cytek Biosciences, USA) and analyzed using FlowJo 10.3 (TreeStar Inc, USA) after sequential doublet exclusion (FSC-A vs. FSC-H and SSC-A vs. SSC-W) and live/dead cell discrimination.

### Lectin stainings

For analysis of lectin binding by immunofluorescence, frozen sections of OCT-embedded tumors were cut and prepared using a cryostat. Biotinylated PNA, MAL II and ConA lectins (Vector Laboratories, USA) were used at 10 µg/mL and incubated in FACS buffer for 20 min at 4°C. Binding of lectins was then detected by incubation with Streptavidin-Cy3, quantified and normalized to the respective area of DAPI binding. For flow cytometric analysis of lectin binding, single cell suspensions of tumor digests were blocked, live/dead stained and incubated with the biotinylated lectins at 10 µg/mL. After detection using Streptavidin-PE-Cy7, samples were fixed (IC fixation buffer, eBioscience, USA) and acquired on a CytoFLEX flow cytometer (Beckman Coulter, USA). Lectin staining was quantified after doublet and live/dead exclusion using the geometric mean fluorescence intensity (MFI). ConA was used as a sialic acid-independent control.

### Multiplex cytokine measurements

Cytokine levels in the serum of B16D5-HER2 bearing C57BL/6 mice were measured 7 days after treatment with E-301, E-301 LOF and trastuzumab. Blood was collected retro-orbitally at the time of sacrifice and clotted for a minimum of 30 min in Microvette 200 Z-Gel tubes containing a clotting activator (Sarstedt, Germany). Clotted samples were centrifuged at 10,000 g for 10 min at room temperature and frozen at −80°C. Measurement of cytokine levels was performed using the Luminex Cytokine & Chemokine 36-Plex Mouse ProcartaPlex™ Panel 1A kit (Thermo Fisher Scientific, USA). Statistical significance was determined using multiple unpaired t-tests with post hoc Bonferroni’s correction for multiple comparisons.

### Generation of sialylation gene expression signatures from TCGA data

First, a set of genes involved in sialic acid biosynthesis and metabolism was generated by merging genes from the Reactome gene set ‘Sialic acid metabolism’ (ID R-HSA-4085001) and the Gene Ontology gene set ‘Sialylation’ (ID GO:0097503). The merged gene set contained the following genes: ST3GAL3, ST6GALNAC5, ST6GALNAC3, ST3GAL5, ST6GAL2, ST3GAL6, ST6GAL1, ST8SIA4, ST3GAL1, ST6GALNAC4, ST6GALNAC6, ST8SIA6, ST3GAL4, ST8SIA1, ST8SIA2, ST3GAL2, ST6GALNAC2, ST6GALNAC1, ST8SIA5, ST8SIA3, C20orf173, NPL, NEU2, NEU4, GLB1, NEU1, SLC17A5, SLC35A1, GNE, NANS, NEU3, CMAS, NANP, and CTSA. The immune gene set used was described previously and contains 3,021 immune-related genes. The expression of each sialylation-associated gene was correlated with the expression of each immune-related gene using the combined TCGA gene expression data of all solid cancers^32^. The correlation matrix was used to perform k-means clustering using k values ranging from 1 to 10. The elbow method indicated that k=5 minimized total intracluster variation, hence a total of five clusters were generated for further analyses. Median expression values of each cluster were used for statistical analyses.

### Survival analysis of TCGA data

For each cancer type, the median of the expression values of the genes in each cluster was calculated and patients divided into quartiles based on their expression values (low, intermediate– low, intermediate–high and high) and the survival analysis was performed for all 5 clusters.

### Correlation with immune cell proportions

The median value of each cluster was correlated with the proportion of 10 immune cell types. The proportions of immune cells were taken from The Cancer Immunome Atlas (TCIA) database (https://tcia.at/cellTypeFractions). The proportions used were retrieved using the method ‘quanTIseq lsei TIL10’.

### Correlation with Immune gene sets from Reactome

Relevant immune gene sets were selected from the Reactome database. Each sialylation gene of a given k-mean cluster was correlated with each gene from the respective immune gene set. The mean of all absolute correlation coefficients was calculated to obtain a correlation score.

### Correlation with T cell dysfunction and exhaustion

The median expression values of gene set 1 from the TCGA datasets LUSC and KIRC were correlated with published T cell dysfunction and exclusion scores^17^.

### Tissue microarray analysis

The tissue microarray HKid-CRCC150CS-01, containing 75 cores of KIRC tumors with adjacent control tissue and survival information was obtained from Biomax (USA). Slides were deparaffinized and stained with a hexameric Siglec-9 Fc construct (Hydra, Palleon, USA) or an anti-CD3 antibody. Binding was detected by IHC, slides scanned and staining intensities quantified using Fiji (ImageJ).

### Patient samples

The local ethics committee in Basel, Switzerland, approved the sample collection and the use of the corresponding clinical data (Ethikkommission Nordwestschweiz, EKNZ, Basel Stadt, Switzerland). Informed consent was obtained from all patients prior to sample collection. Tumor samples were collected locally at the thoracic surgery of the University Hospital Basel, digested and processed as described above and single cell suspensions frozen.

### Human tumor-infiltrating T cell stimulation

Single cell suspensions of human tumor digests were prepared as described above, thawed and seeded in RPMI-1640 medium supplemented with 10% heat-inactivated FBS (PAA Laboratories, Germany), 2 mM L-glutamine, 1 mM sodium pyruvate, 100 µg/mL streptomycin and 100 U/mL penicillin (all Gibco, USA). The cells were then stimulated with 10 ng/mL SEB (Sigma, USA) with or without the addition of 10 mU/mL *Vibrio cholerae* neuraminidase (Roche, Switzerland) for 48 h. Supernatants were frozen at −80°C, and the cells were stained for markers of T cell activation.

### Generation of conditional Siglec-E knockout mice

Siglec-E flox C57BL/6 mice (Elox mice) were made by using a Crispr/Cas9 system (Supplementary Figure 5). A vector carrying 2 loxP sites was used to target the first 3 exons of Siglec-E. CD11-specific conditional Siglec-E deficient animals were made by crossing the CD11-Cre with Siglec-E^loxP/loxP^ mice. Conditional knockout was confirmed by flow cytometry (Figure 5).

### Analysis of scRNA-seq data

Samples were demultiplexed and aligned using Cell Ranger 2.2 (10X genomics) to genome build GRCm38 to obtain a raw read count matrix of barcodes corresponding to cells and features corresponding to detected genes. Read count matrices were processed, analyzed and visualized in R v. 4.0.0 <u>(R Core Team, 2013)</u> using Seurat v. 3^33^ with default parameters in all functions, unless specified. Poor quality cells, with low total unique molecular identifier (UMI) counts and high percent mitochondrial gene expression, were excluded. Filtered samples were normalized using a regularized negative binomial regression (SCTransform)^34^ and integrated with the reciprocal principal component analysis (rpca) approach followed by mutual nearest neighbors, using 50 principal components. Integrated gene expression matrices were visualized with a Uniform Manifold Approximation and Projection (UMAP)^35^ as a dimensionality reduction approach. Resolution for cell clustering was determined by evaluating hierarchical clustering trees at a range of resolutions (0 - 1.2) with Clustree^36^, selecting a value inducing minimal cluster instability. Datasets were subsetted to include only specific cells based on gene expression. Subsetted datasets were then split along conditions, and processed anew as described above.

Differentially expressed genes between clusters were identified as those expressed in at least 25% of cells with a greater than 0.3 natural log fold change and an adjusted p value of less than 0.01, using the FindMarkers function in Seurat v.3 with all other parameters set to default.

### Statistical analysis

Statistical analyses were performed using Prism 9.0 (GraphPad, USA). Comparisons between two groups were performed using the unpaired two-tailed Student’s *t*-test, with the exception of Fig. 1g, where a paired two-tailed *t*-test was used. Differences among more than two groups were assessed using one- or two-way non-parametric analysis of variance (ANOVA), followed by post hoc corrections for multiple comparisons (Tukey’s or Sidak’s). Survival data were analyzed using the log-rank (Mantel–Cox) or the Gehan-Wilcoxon test with post hoc Bonferroni’s correction for multiple comparisons and the multivariate analysis of gene set 1 expression on patient survival was performed by Cox proportional hazard analysis. A *P* value < 0.05 was considered statistically significant. * *P* ≤ 0.05, ** *P* ≤ 0.01, *** *P* ≤ 0.001, **** *P* ≤ 0.0001. n indicates the number of biological replicates, all bars within the graphs represent mean values, and the error bars represent standard errors of the mean.

## Contributions

M.A.S., C.B., L.P, and H.L. conceived and planned the project. M.A.S., N.R.M., N.K., J.W., M.P.T., M.A.G., A.P., K.G. and A.K. performed experiments and interpreted the results. B.K. and G.M. performed statistical analysis of the human expression data. D.E.S analyzed the scRNA-sequencing data. K.N., J. B., O. M. T. P. and E. L. P. provided samples and reagents. M.A.S., A.Z. and H.L. wrote the manuscript. All authors critically read and approved the paper.

## Acknowledgments

This work was supported by funding from the Goldschmidt-Jacobson Foundation (to H.L.), Promedica Foundation (to M.A.S. and A.Z.), Krebsliga Beider Basel (KLBB, to H.L. and M.A.S.), Lichtenstein Foundation (to H.L.), Schoenemakers-Müller Foundation (to H.L.), Swiss National Science Foundation (SNSF Nr. 310030_184720/1 to H.L.), German Research Foundation (DFG) under Germany’s Excellence Strategy (CIBSS EXC-2189 Project ID 390939984 to D.E.S.), as well as a grant from the National Institutes of Health (no. NIH CA227942 to C.R.B.). Researchers were also supported by the National Science Foundation Graduate Research Fellowship (M.A.G.) and the Stanford ChEM-H Chemistry/Biology Interface Predoctoral Training Program (M.A.G.), as well as the Swiss Government Excellence Scholarship for Foreign Scholars and Artists (N.R.M.) and the Conselho Nacional de Desenvolvimento Científico e Tecnológico (N.R.M).

The results shown here are in part based upon data generated by the TCGA Research Network: https://www.cancer.gov/tcga. We thank Petra Herzig, Reto Ritschard, and Mélanie Buchi for their technical assistance. We thank Johan Fridén for his work on the graphical abstract. We also thank all the patients who allowed us to use their materials and made this work possible.

