## Supplemental Figures for "Targeting cancer glycosylation repolarizes tumor-associated macrophages allowing effective immune checkpoint blockade"

#### Extended Figures

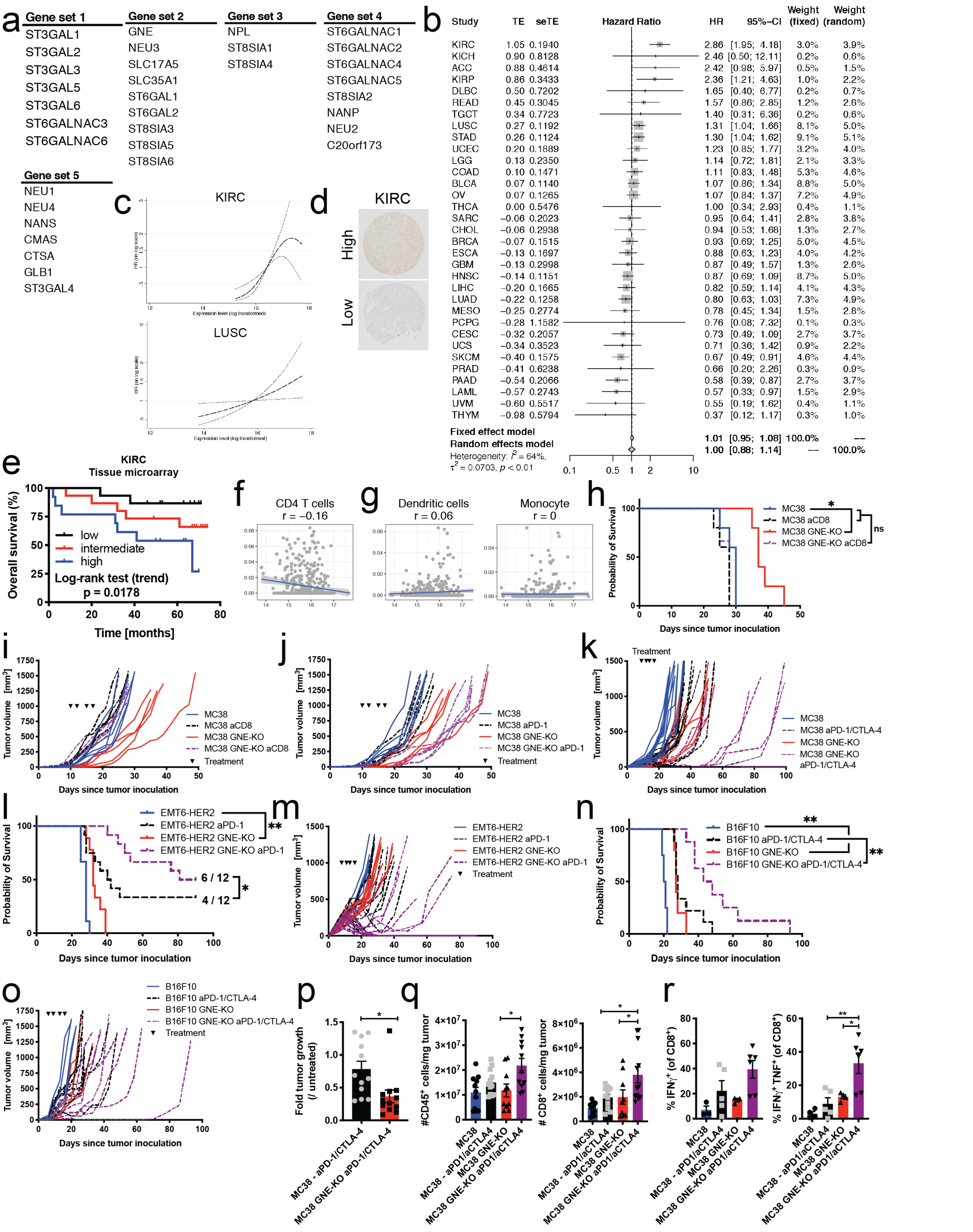
a
Gene set 1
ST3GAL1
ST3GAL2
ST3GAL3
ST3GAL5
ST3GAL6
ST6GALNAC3
ST6GALNAC6
Gene set 2
GNE
NEU3
SLC17A5
SLC35A1
ST6GAL1
ST6GAL2
ST8SIA3
ST8SIA5
ST8SIA6
Gene set 3
NPL
ST8SIA1
ST8SIA4
Gene set 4
ST6GALNAC1
ST6GALNAC2
ST6GALNAC4
ST6GALNAC5
ST8SIA2
NANP
NEU2
C20orf173
Gene set 5
NEU1
NEU4
NANS
CMAS
CTSA
GLB1
ST3GAL4
b
Study TE seTE Hazard Ratio HR 95%-CI Weight (fixed) Weight (random)
KIRC 1.05 0.1940 2.86 [1.95; 4.18] 3.0% 3.9%
KICH 0.90 0.8128 2.46 [0.50; 12.11] 0.2% 0.6%
ACC 0.88 0.4614 2.42 [0.98; 5.97] 0.5% 1.5%
KIRP 0.86 0.3433 2.36 [1.21; 4.63] 1.0% 2.2%
DLBC 0.50 0.7202 1.65 [0.40; 6.77] 0.2% 0.7%
READ 0.45 0.3045 1.57 [0.86; 2.85] 1.2% 2.6%
TGCT 0.34 0.7723 1.40 [0.31; 6.36] 0.2% 0.6%
LUSC 0.27 0.1192 1.31 [1.04; 1.66] 8.1% 5.0%
STAD 0.26 0.1124 1.30 [1.04; 1.62] 9.1% 5.1%
UCEC 0.20 0.1889 1.23 [0.85; 1.77] 3.2% 4.0%
LGG 0.13 0.2350 1.14 [0.72; 1.81] 2.1% 3.3%
COAD 0.10 0.1471 1.11 [0.83; 1.48] 5.3% 4.6%
BLCA 0.07 0.1140 1.07 [0.86; 1.34] 8.8% 5.0%
OV 0.07 0.1265 1.07 [0.84; 1.37] 7.2% 4.9%
THCA 0.00 0.5476 1.00 [0.34; 2.93] 0.4% 1.1%
SARC −0.06 0.2023 0.95 [0.64; 1.41] 2.8% 3.8%
CHOL −0.06 0.2938 0.94 [0.53; 1.68] 1.3% 2.7%
BRCA −0.07 0.1515 0.93 [0.69; 1.25] 5.0% 4.5%
ESCA −0.13 0.1697 0.88 [0.63; 1.23] 4.0% 4.2%
GBM −0.13 0.2998 0.87 [0.49; 1.57] 1.3% 2.6%
HNSC −0.14 0.1151 0.87 [0.69; 1.09] 8.7% 5.0%
LIHC −0.20 0.1665 0.82 [0.59; 1.14] 4.1% 4.3%
LUAD −0.22 0.1258 0.80 [0.63; 1.03] 7.3% 4.9%
MESO −0.25 0.2774 0.78 [0.45; 1.34] 1.5% 2.8%
PCPG −0.28 1.1582 0.76 [0.08; 7.32] 0.1% 0.3%
CESC −0.32 0.2057 0.73 [0.49; 1.09] 2.7% 3.7%
UCS −0.34 0.3523 0.71 [0.36; 1.42] 0.9% 2.2%
SKCM −0.40 0.1575 0.67 [0.49; 0.91] 4.6% 4.4%
PRAD −0.41 0.6238 0.66 [0.20; 2.26] 0.3% 0.9%
PAAD −0.54 0.2066 0.58 [0.39; 0.87] 2.7% 3.7%
LAML −0.57 0.2743 0.57 [0.33; 0.97] 1.5% 2.9%
UVM −0.60 0.5517 0.55 [0.19; 1.62] 0.4% 1.1%
THYM −0.98 0.5794 0.37 [0.12; 1.17] 0.3% 1.0%
Fixed effect model 1.01 [0.95; 1.08] 100.0% --
Random effects model 1.00 [0.88; 1.14] -- 100.0%
Heterogeneity: $I^2$ = 64%, $\tau^2$ = 0.0703, p < 0.01
0.1 0.5 1 2 10
c
KIRC
LUSC
Expression level (log transformed)
d
KIRC
High
Low
e
KIRC
Tissue microarray
Overall survival (%)
low
intermediate
high
Log-rank test (trend)
p = 0.0178
Time [months]
f
CD4 T cells
r = −0.16
g
Dendritic cells
r = 0.06
Monocyte
r = 0
h
Probability of Survival
MC38
MC38 aCD8
MC38 GNE-KO
MC38 GNE-KO aCD8
ns
Days since tumor inoculation
i
Tumor volume [mm³]
MC38
MC38 aCD8
MC38 GNE-KO
MC38 GNE-KO aCD8
Treatment
Days since tumor inoculation
j
Tumor volume [mm³]
MC38
MC38 aPD-1
MC38 GNE-KO
MC38 GNE-KO aPD-1
Treatment
Days since tumor inoculation
k
Treatment
Tumor volume [mm³]
MC38
MC38 aPD-1/CTLA-4
MC38 GNE-KO
MC38 GNE-KO aPD-1/CTLA-4
Days since tumor inoculation
l
Probability of Survival
EMT6-HER2
EMT6-HER2 aPD-1
EMT6-HER2 GNE-KO
EMT6-HER2 GNE-KO aPD-1
6 / 12
4 / 12
Days since tumor inoculation
m
Tumor volume [mm³]
EMT6-HER2
EMT6-HER2 aPD-1
EMT6-HER2 GNE-KO
EMT6-HER2 GNE-KO aPD-1
Treatment
Days since tumor inoculation
n
Probability of Survival
B16F10
B16F10 aPD-1/CTLA-4
B16F10 GNE-KO
B16F10 GNE-KO aPD-1/CTLA-4
Days since tumor inoculation
o
Tumor volume [mm³]
B16F10
B16F10 aPD-1/CTLA-4
B16F10 GNE-KO
B16F10 GNE-KO aPD-1/CTLA-4
Treatment
Days since tumor inoculation
p
Fold tumor growth (/ untreated)
MC38 - aPD-1/CTLA-4
MC38 GNE-KO aPD-1/CTLA-4
q
#CD45⁺ cells/mg tumor
### CD8⁺ cells/mg tumor
MC38
MC38 - aPD1/aCTLA4
MC38 GNE-KO
MC38 GNE-KO aPD1/aCTLA4
r
% IFNγ⁺ (of CD8⁺)
% IFNγ⁺ TNF⁺ (of CD8⁺)

**Extended Data Figure 1. Tumor sialylation is associated with immune suppression and reduced survival in cancer patients.**

**a**, Gene sets 1–5 generated from the clustering of genes involved in sialic acid biosynthesis with immune genes using all solid cancers in TCGA database. **b**, Forest tree plot of the hazard ratios of increasing gene set 1 expression for all solid cancer types in the TCGA. **c**, Plot of continuous gene set 1 expression and the corresponding hazard rations in clear cell renal cell carcinoma (KIRC) and squamous cell carcinoma of the lung (LUSC). **d**, Representative KIRC tissue cores of tumors with high and low intensities of Siglec-9 Fc staining, from a tissue microarray of 75 KIRC patients**. e**, Kaplan–Meier survival curve of KIRC patients, divided into terciles based on the intensity of Siglec-9 Fc staining. **f**, Correlation of gene set 1 expression with a gene expression signature of conventional $CD4^{+}$ T cells, for all LUSC patients in the TCGA database. **g**, Correlation of gene set 1 expression with a gene expression signature of dendritic cells (DCs) and monocytes, for all LUSC patients in the TCGA database. **h**, Impact of $CD8^{+}$ T cell depletion on the survival of mice carrying MC38 wildtype and GNE-KO tumors. **i**, Impact of $CD8^{+}$ T cell depletion on the growth of individual MC38 wildtype and GNE-KO tumors. **j**, Effect of anti-PD-1 ICB on the growth of individual MC38 wildtype and GNE-KO tumors. **k**, Effect of anti-PD-1 and anti-CTLA-4 ICB on the growth of individual MC38 wildtype and GNE-KO tumors. **l**, Effect of PD-1 blockade on the survival of mice bearing intramammary wildtype or GNE-KO EMT6-HER2 tumors (n=9–12 mice per group). **m**, Effect of an anti-PD-1 ICB on the growth of individual EMT6-HER2 wildtype and GNE-KO tumors. **n**, Effect of combined PD-1 and CTLA-4 blockade on the survival of mice bearing subcutaneous wildtype or GNE-KO B16F10 tumors (n=4–8 mice per group). **o**, Effect of anti-PD-1 and anti-CTLA-4 ICB on the growth of individual B16F10 wildtype and GNE-KO tumors. **p**, Relative tumor growth over treatment period, compared to untreated. **q**, Absolute number of $CD45^{+}$ immune cells and $CD8^{+}$ T cells per mg of resected tumor. **f**, Frequency of $IFN\gamma^{+}$ and $IFN\gamma^{+}TNF^{+}$ $CD8^{+}$ T cells after *ex vivo* PMA/ionomycin restimulation. n indicates the number of biological replicates. Error bars represent the mean ± standard error of the mean (s.e.m.). Statistical analyses were performed using the log-rank (Mantel–Cox) test for the TCGA survival data or the Gehan-Wilcoxon test for the mouse survival data, followed by Bonferroni's correction for multiple comparisons. An unpaired two-tailed Student's *t*-test was used in 1p and one-way ANOVAs with post hoc Sidak's test 1q, r. * $P \leq 0.05$, ** $P \leq 0.01$, *** $P \leq 0.001$, **** $P \leq 0.0001$.

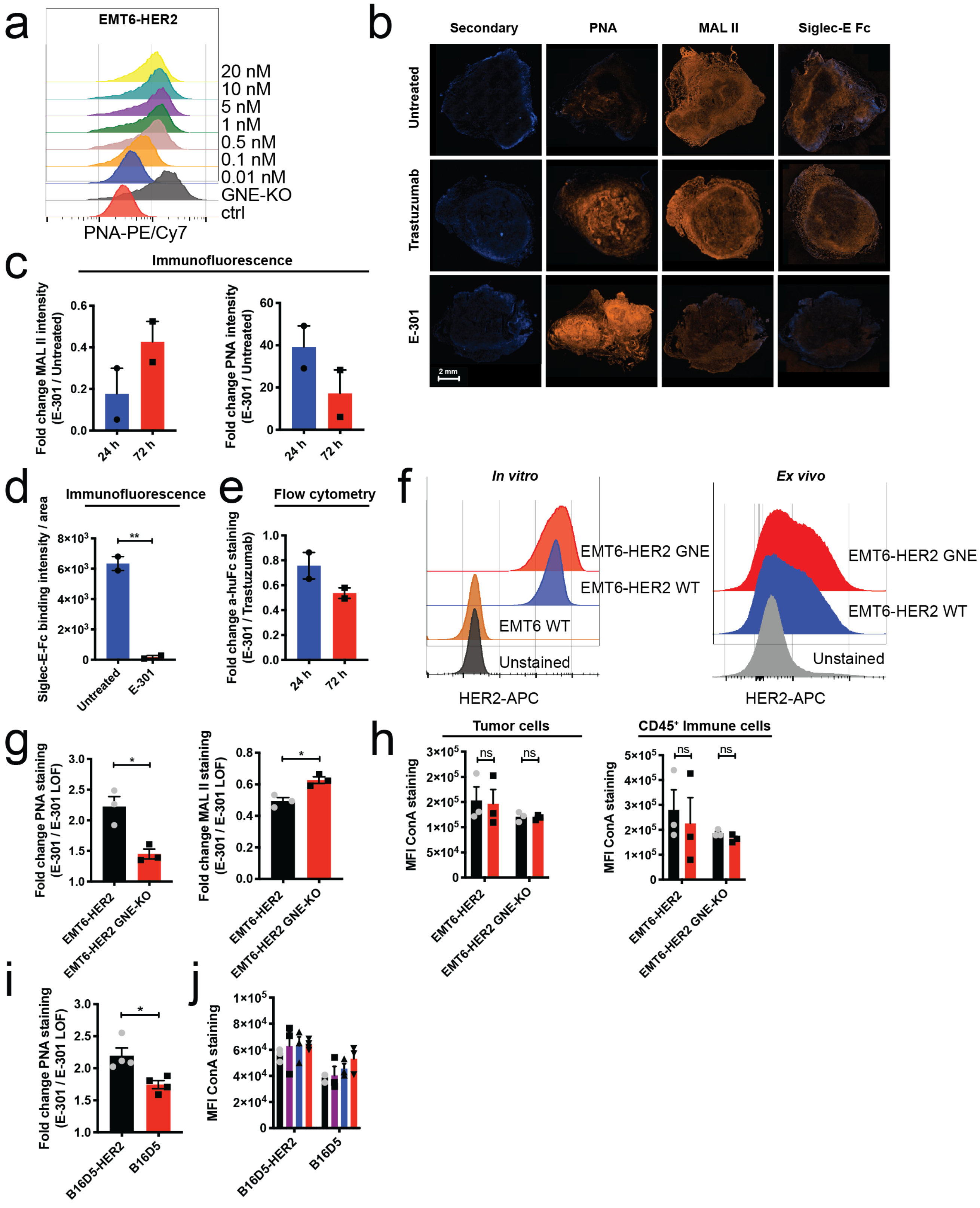
a
EMT6-HER2
20 nM
10 nM
5 nM
1 nM
0.5 nM
0.1 nM
0.01 nM
GNE-KO
ctrl
PNA-PE/Cy7
b
Secondary
PNA
MAL II
Siglec-E Fc
Untreated
Trastuzumab
E-301
2 mm
c
Immunofluorescence
Fold change MAL II intensity (E-301 / Untreated)
Fold change PNA intensity (E-301 / Untreated)
24 h
72 h
d
Immunofluorescence
Siglec-E-Fc binding intensity / area
Untreated
E-301
e
Flow cytometry
Fold change a-huFc staining (E-301 / Trastuzumab)
f
In vitro
Ex vivo
EMT6-HER2 GNE
EMT6-HER2 WT
EMT6 WT
Unstained
HER2-APC
g
Fold change PNA staining (E-301 / E-301 LOF)
Fold change MAL II staining (E-301 / E-301 LOF)
EMT6-HER2
EMT6-HER2 GNE-KO
h
Tumor cells
CD45⁺ Immune cells
MFI ConA staining
ns
i
B16D5-HER2
B16D5
j

**Extended Data Figure 2. Tumor-targeted sialidase effectively desialylates the tumor microenvironment.**

**a**, Representative histograms of PNA staining after 24 h *in vitro* incubation of EMT6-HER2 cells with increasing concentrations of trastuzumab, E-301 LOF or E-301. EMT6-HER2 GNE-KO cells were used as a control for desialylation. **b**, Representative lectin and Siglec-E Fc stained immunofluorescence images of untreated, trastuzumab- or E-301-treated EMT6-HER2 tumors at 72 h post-treatment. **c**, Fold changes in MAL II and PNA staining intensities relative to those of the untreated control. **d**, Quantification of immunofluorescence staining of Siglec-E Fc (72 h). The sum of the staining intensity was normalized to the respective DAPI-stained area. **e**, Fold change in the MFI of secondary anti-human Fc staining 24 h and 72 h after E-301 treatment relative to that after trastuzumab treatment (b–e, n=2 mice per group). **f**, anti-HER2 staining of wildtype EMT6-HER2 and EMT6-HER2 GNE-KO tumor cells, after *in vitro* culture and isolated *ex vivo* from established tumors. **g**, Fold changes in PNA and MAL II staining intensities of E-301 treatment relative to E-301 LOF treated control tumors, both for wildtype EMT6-HER2 and EMT6-HER GNE-KO tumors. **h**, MFI of ConA staining after E-301- or E-301 LOF-treatment of wildtype EMT6-HER2 and EMT6-HER2 GNE-KO tumors, both for tumors cells and tumor-infiltrating $CD45^+$ immune cells. **i**, Fold change in the geometric MFI of PNA staining after E-301 treatment relative to E-301 LOF treatment (n=4). **j**, MFI ConA staining of untreated-, trastuzumab-, E-301 LOF-or E-301-treated tumors. n indicates the number of biological replicates. Error bars represent the mean ± standard error of the mean (s.e.m.). Statistical analyses were performed using unpaired two-tailed Student's *t*-test , or one- and two-way ANOVAs with post hoc Sidak's test for group comparisons. * $P \leq 0.05$, ** $P \leq 0.01$, *** $P \leq 0.001$, **** $P \leq 0.0001$.

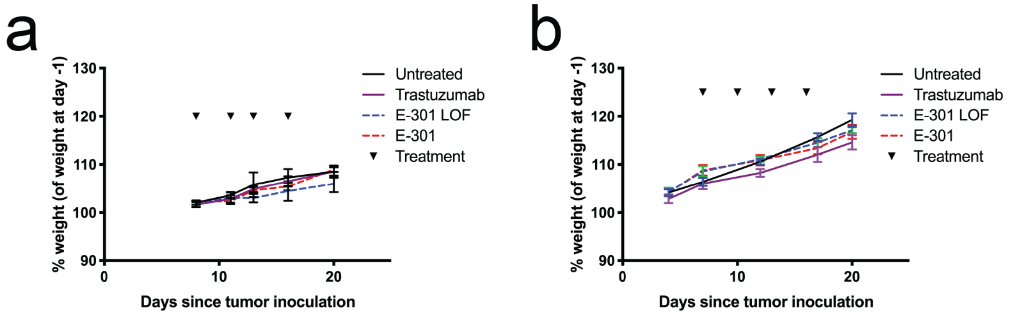


**Extended Data Figure 3. Tumor-targeted sialidase inhibits tumor growth by activating the adaptive immune system.**

**a**, Body weights of mice bearing intramammary EMT6-HER2 tumors after treatment with trastuzumab, E-301 LOF or E-301 relative to weight before first treatment. **b**, Body weights of mice bearing subcutaneous B16D5-HER2 tumors after treatment with trastuzumab, E-301 LOF or E-301 relative to before treatment. Error bars represent the mean ± s.e.m.

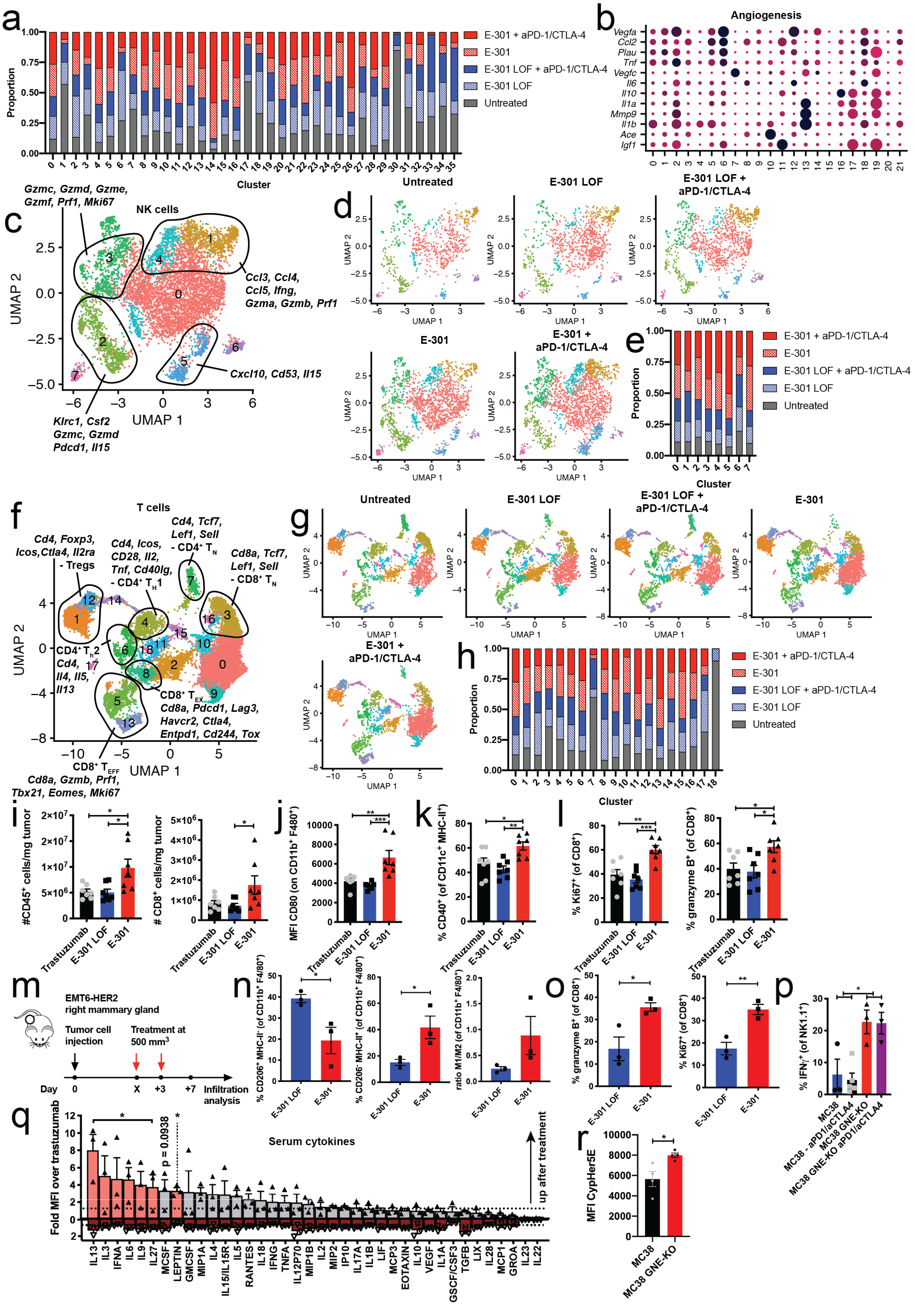
a
1.00
0.75
0.50
0.25
0.00
Proportion
Cluster
E-301 + aPD-1/CTLA-4
E-301
E-301 LOF + aPD-1/CTLA-4
E-301 LOF
Untreated
b
Angiogenesis
Vegfa
Ccl2
Plau
Tnf
Vegfc
Il6
Il10
Il1a
Mmp9
Il1b
Ace
Igf1
c
NK cells
Gzmc, Gzmd, Gzme, Gzmf, Prf1, Mki67
Ccl3, Ccl4, Ccl5, Ifng, Gzma, Gzmb, Prf1
Cxcl10, Cd53, Il15
Klrc1, Csf2 Gzmc, Gzmd Pdcd1, Il15
UMAP 1
UMAP 2
d
Untreated
E-301 LOF
E-301 LOF + aPD-1/CTLA-4
E-301
E-301 + aPD-1/CTLA-4
e
f
T cells
Cd4, Foxp3, Icos,Ctla4, Il2ra - Tregs
Cd4, Icos, CD28, Il2, Tnf, Cd40lg, - CD4+ TH1
Cd4, Tcf7, Lef1, Sell - CD4+ TN
Cd8a, Tcf7, Lef1, Sell - CD8+ TN
CD4+ TH2 Cd4, Il4, Il5, Il13
CD8+ TEX Cd8a, Pdcd1, Lag3, Havcr2, Ctla4, Entpd1, Cd244, Tox
CD8+ TEFF Cd8a, Gzmb, Prf1, Tbx21, Eomes, Mki67
g
h
i
#CD45+ cells/mg tumor
### CD8+ cells/mg tumor
Trastuzumab
E-301 LOF
E-301
j
MFI CD80 (on CD11b+ F480+)
k
% CD40+ (of CD11c+ MHC-II+)
l
% Ki67+ (of CD8+)
% granzyme B+ (of CD8+)
m
EMT6-HER2 right mammary gland
Tumor cell injection
Treatment at 500 mm3
Day
Infiltration analysis
n
% CD206+ MHC-II- (of CD11b+ F4/80+)
% CD206- MHC-II+ (of CD11b+ F4/80+)
ratio M1/M2 (of CD11b+ F4/80+)
o
p
% IFNγ+ (of NK1.1+)
MC38
MC38 - aPD1/aCTLA4
MC38 GNE-KO
MC38 GNE-KO aPD1/aCTLA4
q
Fold MFI over trastuzumab
Serum cytokines
p = 0.0938
up after treatment
IL13
IL3
IFNA
IL6
IL9
IL27
MCSF
LEPTIN
GMCSF
MIP1A
IL4
IL15/IL15R
IL5
RANTES
IL18
IFNG
TNFA
IL12P70
MIP1B
IL2
MIP2
IP10
IL17A
IL1B
LIF
MCP3
EOTAXIN
IL10
VEGF
IL1A
GSCF/CSF3
TGFB
LIX
IL28
MCP1
GROA
IL23
IL22
r
MFI CypHer5E
MC38
MC38 GNE-KO

**Extended Data Figure 4. Tumor-targeted desialylation repolarizes tumor-associated macrophages.** **a**, Contribution of each condition to each cluster of $CD45^+$ cells. **b**, Dot plot representation of differentially expressed genes between the macrophage clusters. Size reflects the percentage of each cluster expressing a given gene, average scaled expression is indicated on the color gradient. **c**, Subclustering and UMAP projection of all NK cells. **d**, UMAP projections of NK cells are shown separated by condition. **e**, Contribution of each condition to each NK cell cluster. **f**, Subclustering and UMAP projection of all T cells. **g**, UMAP projections of T cells are shown separated by condition. **h**, Contribution of each condition to each T cell cluster. **i**, Absolute number of total $C45^+$ and $CD8^+$ T cells per mg of resected tumor. **j**, MFI of CD80 expression on $CD11b^+F4/80^+$ TAMs. **k**, Frequency of CD40 expression on DCs. **l**, Frequency of granzyme B and Ki67 expression on $CD8^+$ T cells (i–l, n=7). **m**, Experimental setup for immunophenotyping analysis of changes in immune infiltrates after E-301 treatment: Mice bearing established (500 $mm^3$) subcutaneous EMT6-HER2 tumors were treated with two doses of 10 mg/kg trastuzumab, E-301 LOF or E-301 and immune infiltrates analyzed after 7 days by flow cytometry. **n**, Frequencies of $CD206^-MHC\text{-}II^+$ (M1) and $CD206^+MHC\text{-}II^-$ (M2) cells among $CD11b^+F4/80^+$ tumor-associated macrophages. Ratio of M1 to M2 macrophages in $CD11b^+F4/80^+$ cells. **o**, Expression of granzyme B and Ki67 by $CD8^+$ T cells (n, o, n=3). **p**, Frequency of $IFN\gamma^+$ NK cells after *ex vivo* PMA/ionomycin restimulation (n=3–5). **q**, Luminex analysis of cytokine levels in the serum of mice bearing subcutaneous B16D5-HER2 tumors treated with E-301, E-301 LOF and trastuzumab at day 7 (n=3). Red bars of E-301 samples represent significant changes ($p < 0.05$). **r**, Quantification of phagocytosis measured by CypHer5E intensity in $CD11b^+F4/80^+$ peritoneal macrophages after i.p. injection of CypHer5E labelled wildtype and GNE-KO MC38 tumor cells (n=3). n indicates the number of biological replicates. Error bars represent the mean ± standard error of the mean (s.e.m.). Statistical analyses were performed using one-way ANOVAs with post hoc Tukey's test or unpaired two-tailed Student's *t*-test to assess differences between two groups. * $P \leq 0.05$, ** $P \leq 0.01$, *** $P \leq 0.001$, **** $P \leq 0.0001$.

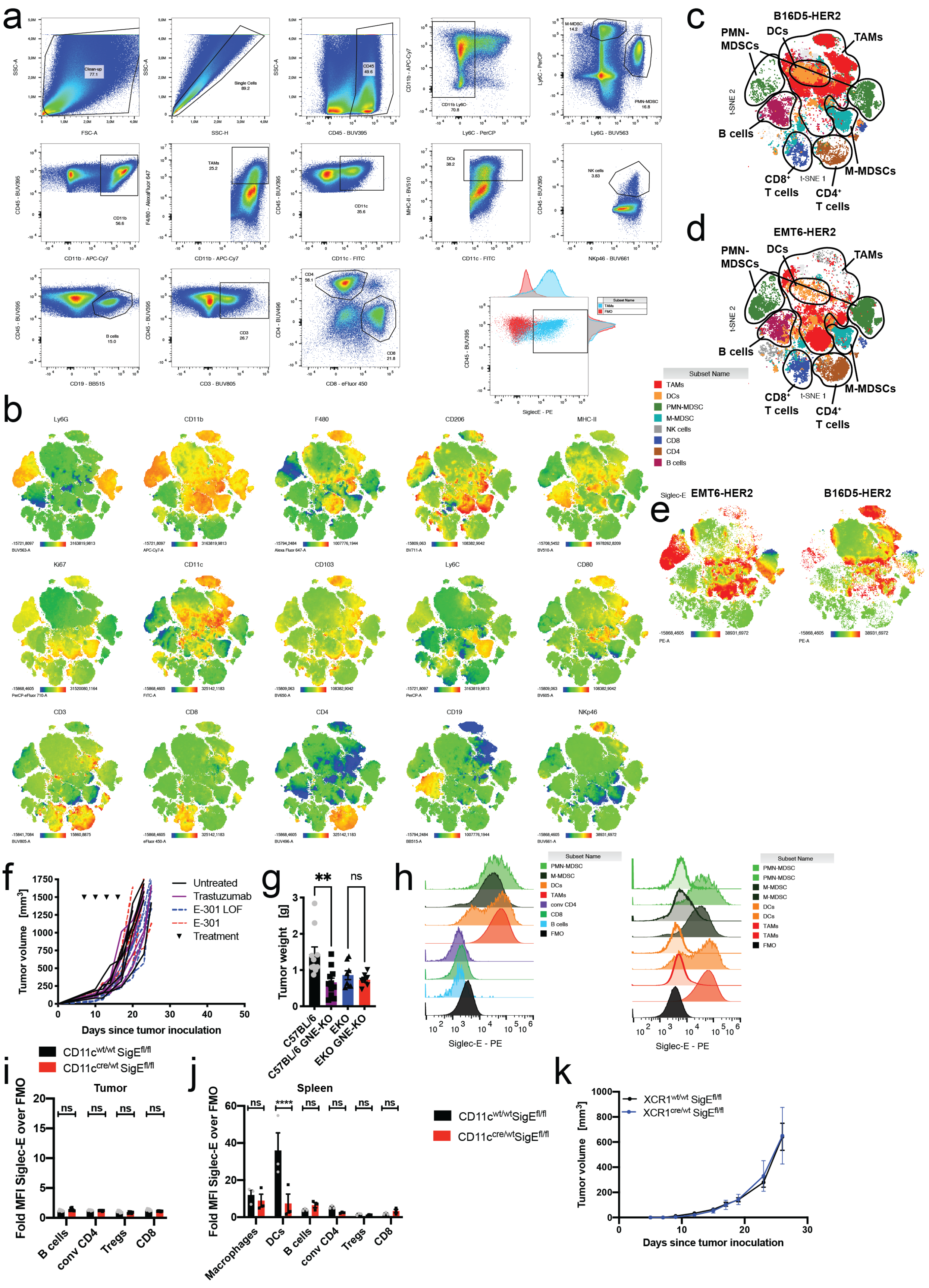
a
b
c
B16D5-HER2
PMN-MDSCs
DCs
TAMs
B cells
CD8+ T cells
CD4+ T cells
M-MDSCs
t-SNE 1
t-SNE 2
d
EMT6-HER2
Subset Name
TAMs
DCs
PMN-MDSC
M-MDSC
NK cells
CD8
CD4
B cells
e
Siglec-E
EMT6-HER2
B16D5-HER2
Ly6G
CD11b
F480
CD206
MHC-II
Ki67
CD11c
CD103
Ly6C
CD80
CD3
CD8
CD4
CD19
NKp46
f
Tumor volume [mm3]
Days since tumor inoculation
Untreated
Trastuzumab
E-301 LOF
E-301
Treatment
g
Tumor weight [g]
C57BL/6
C57BL/6 GNE-KO
EKO
EKO GNE-KO
ns
h
Siglec-E - PE
conv CD4
FMO
i
CD11cwt/wt SigEfl/fl
CD11ccre/wt SigEfl/fl
Tumor
Fold MFI Siglec-E over FMO
B cells
conv CD4
Tregs
CD8
j
Spleen
Macrophages
k
XCR1wt/wt SigEfl/fl
XCR1cre/wt SigEfl/fl
Tumor volume [mm3]
Days since tumor inoculation

**Extended Data Figure 5. Efficacy of tumor-targeted sialidase is dependent of Siglec-E on TAMs.** **a**, Exemplary gating strategy used for Fig. 5c. **b**, t-distributed stochastic neighbor embedding (t-SNE) projection of multicolor flow cytometric immunophenotyping of B16D5-HER2 and EMT6-HER2 tumors. Expression of individual markers is indicated as color gradients from blue (low) to red (high). **c**, t-SNE projection of multicolor flow cytometric immunophenotyping of B16D5-HER2 tumors. Cell populations have been assigned based on maker expression. **d**, t-SNE projection of multicolor flow cytometric immunophenotyping of EMT6-HER2 tumors. Cell populations have been assigned based on maker expression (d, e, 5 concatenated tumor samples each). **e**, Staining intensity for Siglec-E is shown as a color gradient from blue (low) to red (high). **f**, Tumor growth in EKO mice bearing subcutaneous B16D5-HER2 tumors after trastuzumab, E-301 LOF or E-301 treatment (n=6 mice per group). **g**, Tumor weights of MC38 wildtype and GNE-KO tumors from wildtype C57BL/6 and EKO mice (n=13–17 mice per group). **h**, Representative flow cytometry histograms of anti-Siglec-E stainings of single cell suspensions of MC38 tumors from Elox mice crossed to CD11c-Cre mice. **i**, Siglec-E expression by flow cytometry on different lymphoid tumor-infiltrating immune cells in CD11c-Cre Elox mice compared to littermate control mice. Siglec-E expression shown as fold change over control staining. **j**, Siglec-E expression by flow cytometry on different splenic immune cells in CD11c-Cre Elox mice compared to littermate control mice. Siglec-E expression shown as fold change over control staining. **k**, Tumor growth of subcutaneously injected MC38 cells in XCR1-Cre mice crossed to Siglec-E flox (Elox) mice or littermate control mice (Elox only, n=7–8). n indicates the number of biological replicates. Error bars represent the mean ± standard error of the mean (s.e.m.). Statistical analyses were performed using two-way ANOVAs with post hoc Tukey's test. * $P \leq 0.05$, ** $P \leq 0.01$, *** $P \leq 0.001$, **** $P \leq 0.0001$.

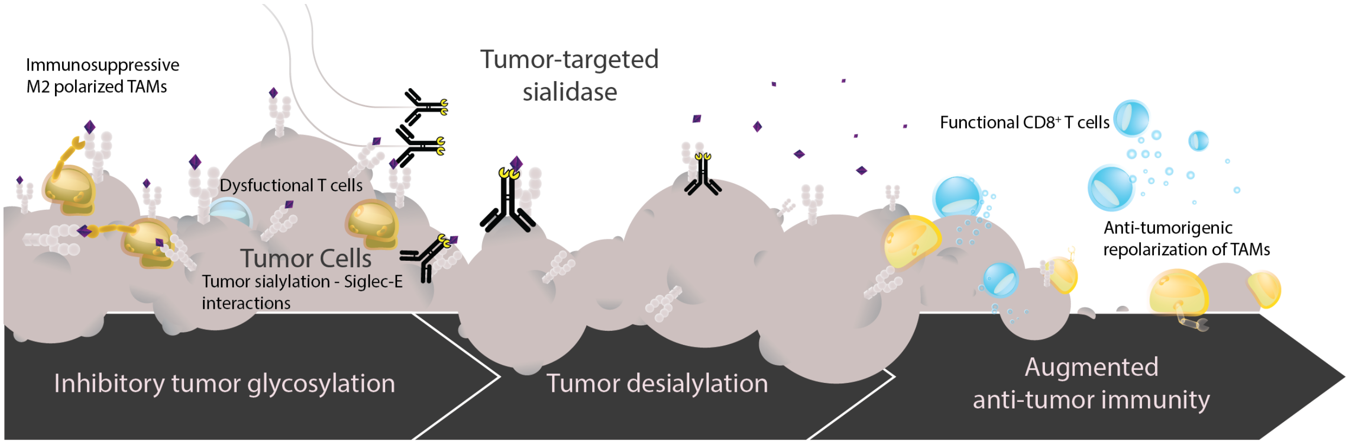


**Extended Data Figure 6. Graphical abstract.**
